# Replication Fork Slowing and Stalling are Distinct, Checkpoint-Independent Consequences of Replicating Damaged DNA

**DOI:** 10.1101/122895

**Authors:** Divya Ramalingam Iyer, Nicholas Rhind

**Affiliations:** Department of Biochemistry & Molecular Pharmacology, University of Massachusetts Medical School, Worcester, Massachusetts, United States of America

## Abstract

In response to DNA damage during S phase, cells slow DNA replication. This slowing is orchestrated by the intra-S checkpoint and involves inhibition of origin firing and reduction of replication fork speed. Slowing of replication allows for tolerance of DNA damage and suppresses genomic instability. Although the mechanisms of origin inhibition by the intra-S checkpoint are understood, major questions remain about how the checkpoint regulates replication forks: Does the checkpoint regulate the rate of fork progression? Does the checkpoint affect all forks, or only those encountering damage? Does the checkpoint facilitate the replication of polymerase-blocking lesions? To address these questions, we have analyzed the checkpoint in the fission yeast *Schizosaccharomyces pombe* using a single-molecule DNA combing assay, which allows us to unambiguously separate the contribution of origin and fork regulation towards replication slowing, and allows us to investigate the behavior of individual forks. Moreover, we have interrogated the role of forks interacting with individual sites of damage by using three damaging agents—MMS, 4NQO and bleomycin—that cause similar levels of replication slowing with very different frequency of DNA lesions. We find that the checkpoint slows replication by inhibiting origin firing, but not by decreasing fork rates. However, the checkpoint appears to facilitate replication of damaged templates, allowing forks to more quickly pass lesions. Finally, using a novel analytic approach, we rigorously identify fork stalling events in our combing data and show that they play a previously unappreciated role in shaping replication kinetics in response to DNA damage.

**Author Summary:** Faithful duplication of the genome is essential for genetic stability of organisms and species. To ensure faithful duplication, cells must be able to replicate damaged DNA. To do so, they employ checkpoints that regulate replication in response to DNA damage. However, the mechanisms by which checkpoints regulate DNA replication forks, the macromolecular machines that contain the helicases and polymerases required to unwind and copy the parental DNA, is unknown. We have used DNA combing, a single-molecule technique that allows us to monitor the progression of individual replication forks, to characterize the response of fission yeast replication forks to DNA damage that blocks the replicative polymerases. We find that forks pass most lesions with only a brief pause and that this lesion bypass is checkpoint independent. However, at a low frequency, forks stall at lesions, and that the checkpoint is required to prevent these stalls from accumulating single-stranded DNA. Our results suggest that the major role of the checkpoint is not to regulate the interaction of replication forks with DNA damage, *per se*, but to mitigate the consequences of fork stalling when forks are unable to successfully navigate DNA damage on their own.

## Introduction

In response to DNA damage during the G1 or G2 phase of the cell cycle, DNA damage checkpoints block cell cycle progression, giving cells time to repair damage before proceeding to the next phase of the cell cycle (Hartwell and Weinert, 1989; Rhind and Russell, 2012). The response to DNA damage during S phase is more complicated, because repair has to be coordinated with ongoing DNA replication (Bartek et al., 2004). DNA damage during S phase activates the intra-S DNA damage checkpoint, which does not completely block S-phase progression, but rather slows DNA replication, presumably allowing for replication-coupled repair (Rhind and Russell, 2000). Lack of the intra-S DNA damage checkpoint predisposes cells to genomic instability (Zhou and Elledge, 2000). Nonetheless, the mechanisms by which replication is slowed, and the roles of checkpoint-dependent and -independent regulation in the S-phase DNA damage response, are not well understood.

The slowing of S phase in response to damage involves both inhibition of origin firing and reduction in fork rate (Kaufmann et al., 1980; Santocanale and Diffley, 1998; Falck et al., 2002; Merrick et al., 2004; Chastain et al., 2006; Seiler et al., 2007; Kumar and Huberman, 2009). The effect of the checkpoint on origin firing has been characterized in budding yeast and mammalian cells. The checkpoint prevents activation of late origins by targeting initiation factors required for origin firing. In mammals, checkpoint kinase 1 (Chk1) inhibits origin firing by targeting the replication kinases, cyclin-dependent kinase (CDK) and Dbf4-dependent kinase (DDK) (Falck et al., 2001; Zhao and Piwnica-Worms, 2001; Sørensen et al., 2003). In budding yeast, Rad53 targets Sld3, an origin initiation factor, and Dbf4, the regulatory subunit of DDK (Lopez-Mosqueda et al., 2010; Zegerman and Diffley, 2010).

Although checkpoint inhibition of origin firing is conserved from yeast to mammals, the contribution of origin regulation to damage tolerance is not clear. For instance, budding yeast mutants such as *mec 1-100*, *SLD3-m25 and dbf4-m25*, which cannot block origin firing in response to damage, are not sensitive to damaging agents such as methyl methanesulfonate (MMS) (Paciotti et al., 2001; Tercero et al., 2003; Lopez-Mosqueda et al., 2010; Zegerman and Diffley, 2010) suggesting that checkpoint regulation of origin firing is not as critical as the checkpoint’s contribution to damage tolerance via fork regulation.

The effect of checkpoint activation on replication forks is less well understood. Because many DNA damage lesions block the replicative polymerases, for forks to pass leading-strand lesions, they must have some way to reestablish leading-strand synthesis downstream of the lesion. Recruitment of trans-lesion polymerases, template switching and leading-strand repriming have all been proposed to be involved in the fork by-pass of lesions, but which is actually used *in vivo* and how the checkpoint may affect that choice, is unclear (Branzei and Foiani, 2005; Lee and Myung, 2008; Branzei and Foiani, 2009; Daigaku et al., 2010; Sale, 2012; Ulrich, 2012).

A consistent observation is that forks slow in response to DNA damage. Whether slowing of forks in the presence of damage is checkpoint-dependent or simply due to the physical presence of lesions is not clear (Tercero and Diffley, 2001; Unsal-Kaçmaz et al., 2007; Szyjka et al., 2008). Initial work in budding yeast showed that replication forks in checkpoint mutant and wild-type cells were slowed to the same extent in the presence of damage, suggesting that slowing is checkpoint-independent (Tercero and Diffley, 2001). However, subsequent work showed that checkpoint activation inhibited replication of damaged DNA, suggesting an active role in the slowing of replication forks (Szyjka et al., 2008). Furthermore, work in mammalian cells showed a role for checkpoint signaling in DNA-damage-dependent slowing of replication forks (Unsal-Kaçmaz et al., 2007). Checkpoint regulation clearly affects fork stability (Lopes et al., 2001; Cobb et al., 2003; Noguchi et al., 2004; Cobb et al., 2005), but that conclusion is largely based on the response to hydroxyurea (HU)-induced replication stress, which blocks fork progression due to deoxynucleoside triphosphate (dNTP) depletion, and thus does not directly address the role of checkpoint regulation in the replication of damaged templates.

The question of how replication forks are regulated in response to DNA damage is complicated by the difficulty of measuring fork progression rates. To directly address this question requires single-molecule resolution. Bulk assays of replication kinetics, such as the quantitation of radioactive thymidine incorporation or flow cytometry, provide only an average profile of replication kinetics, convolving the effects of origin firing and fork progression and obscuring any heterogeneity in fork slowing. The effects of DNA damage on specific origins and on forks replicating specific loci can be analyzed by gel- or sequence-based methods (Santocanale and Diffley, 1998; Shirahige et al., 1998; Tercero and Diffley, 2001; Szyjka et al., 2008; Kumar and Huberman, 2009), but these techniques still only reveal the average response to DNA damage. Such approaches lack the single-molecule resolution necessary to identify heterogeneity in response to damage and to distinguish, for instance, if all forks pause briefly at all lesions or if only a fraction of forks stall, but for a longer time. Therefore, to investigate the effect of polymerase blocking lesions on individual origins and forks on a global scale, we have used a single-molecule DNA combing assay.

DNA combing is a single-fiber visualization technique that allows mapping of thymidine-analog incorporation patches on uniformly-stretched, megabase-length DNA fibers press; Bensimon et al., 1994; Michalet et al., 1997; Jackson and Pombo, 1998; Bianco et al., 2012; Gallo et al., 2016a; Gallo et al., 2016b). The analysis of replication kinetics by DNA combing involves isolating DNA from cells pulse-labeled *in vivo* with thymidine analogs. Labeled DNA is stretched on a coverslip and DNA replicated during the pulse is identified by immunofluorescence. Sequential labeling with two different analogs allows us to determine the direction and speed of replication of labeled tracks on a fiber (Iyer et al., in press; Jackson and Pombo, 1998) (Figure 1). Because combing reveals the behavior of individual replication forks, it allows us to unambiguously study the effect of checkpoint on origin firing and fork rates. In particular, it allows us to measure the heterogeneity of fork rates and determine if all forks respond the same way to lesions. The importance of measuring the heterogeneity of fork rates can be illustrated in a case where some forks pass lesions without slowing, but others stop at the lesion and do not resume replication for the duration of the experiment, a condition we refer to as fork stalling. The fork stalling is particularly difficult to analyze because stalled forks create ambiguous patterns of nucleotide incorporation which cannot be definitively interpreted in isolation (Técher et al., 2013). However, such stalling events can, in principle, be unambiguously identified on fibers that are long enough to contain multiple replicons, allowing potentially ambiguous incorporation patterns to be definitively interpreted from the context of surrounding forks (Figure 2). We have, for the first time, developed an analysis strategy that rigorously incorporates fork stalling and used it to unambiguously quantitate fork stalling rates and the effect of checkpoint activation on fork stalling.

**Figure 1:**
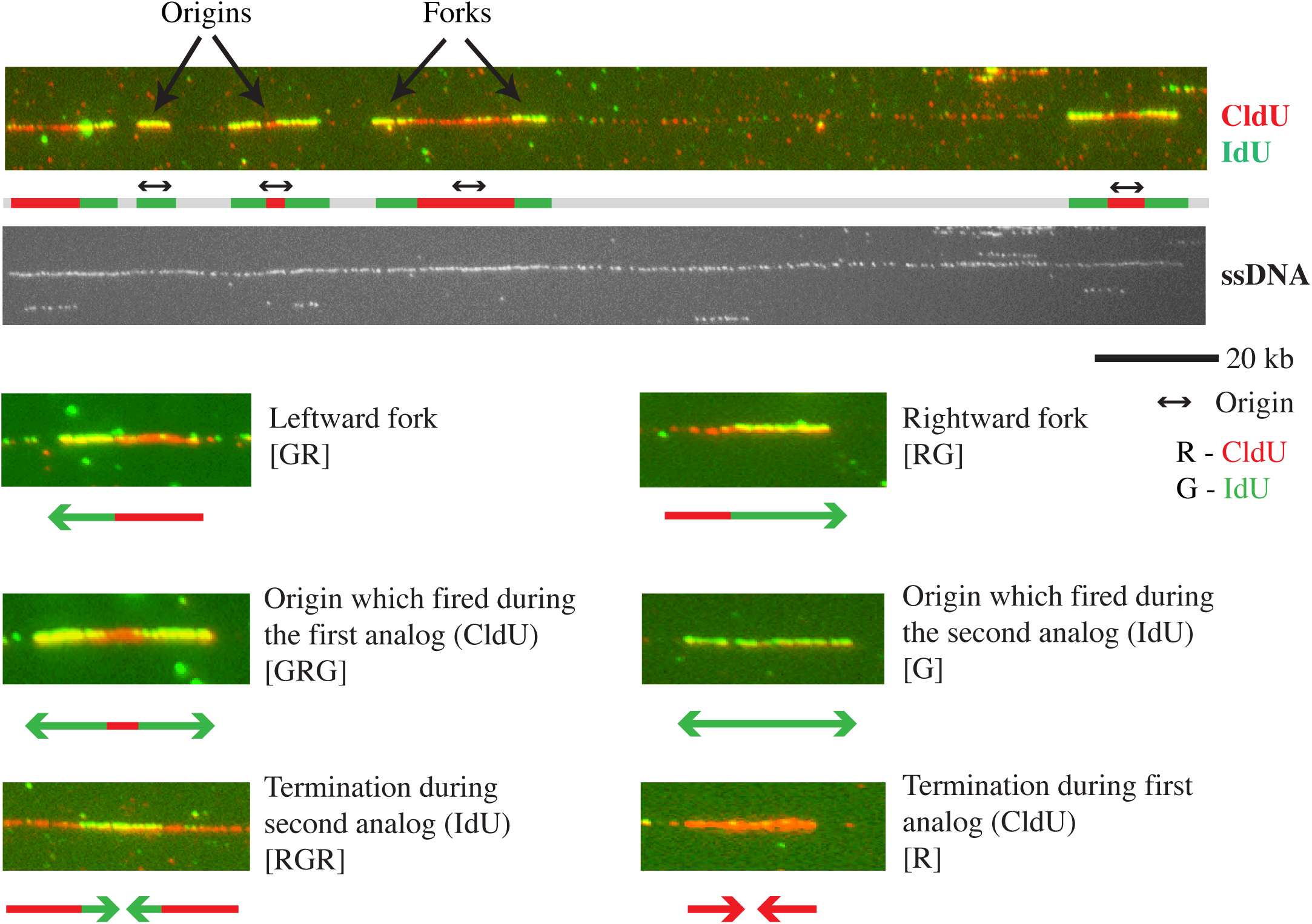
Visualization of replication fork progression. Wild-type cells (yFS940) were G1 synchronized and pulse labeled at 50 minutes after release into S phase with CldU (visualized using red antibody) for 5 minutes, and chased with IdU (visualized using green antibody) for 10 minutes. Shown are a sample fiber from the wild-type untreated dataset and the various replication patterns observed in the dataset with their simplest interpretations. See Figure 2 for other possible interpretations of ambiguous patterns.

**Figure 2:**
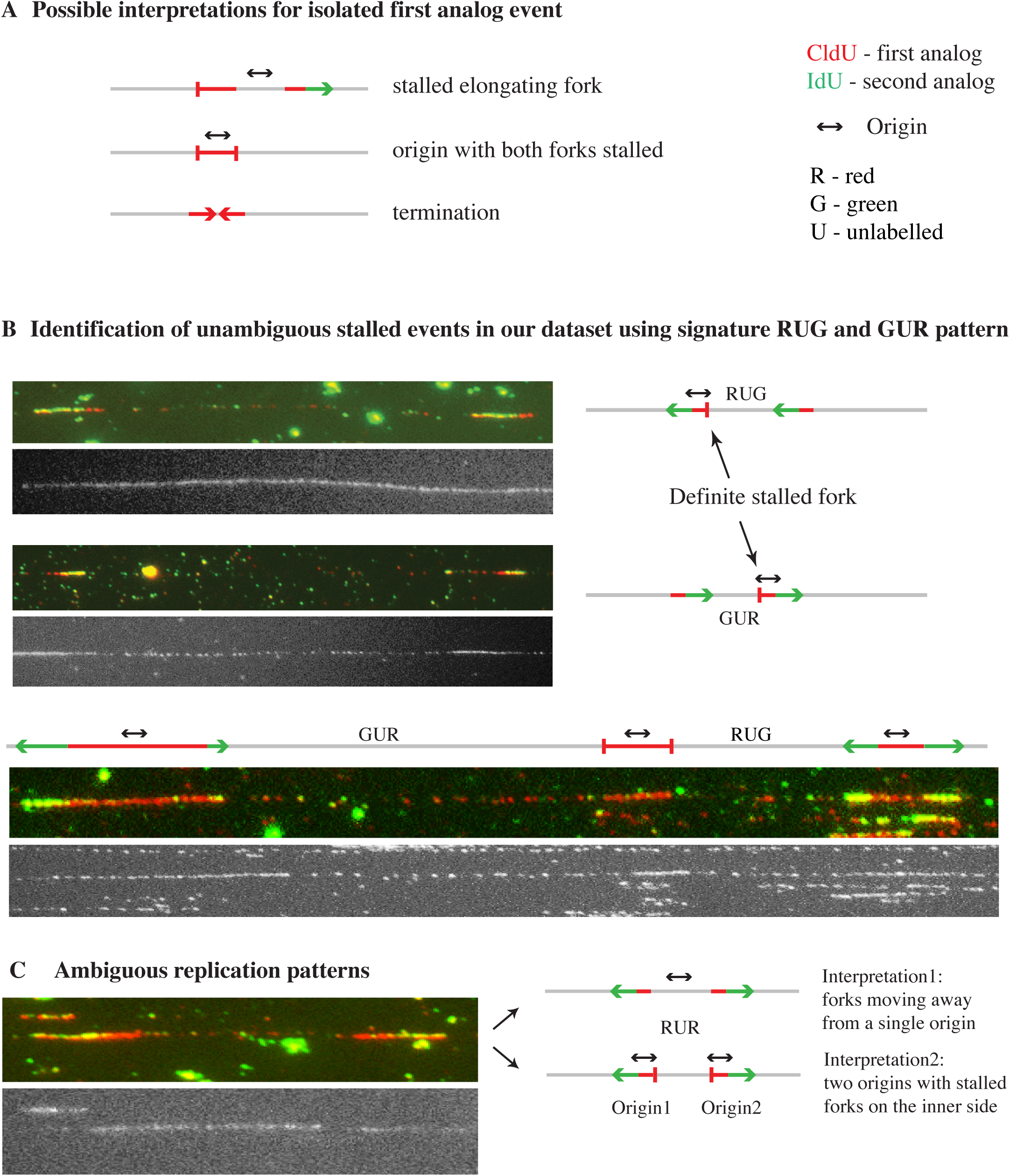
Identification of stalled forks using the context of neighboring forks. (A) Ambiguity in interpreting isolated first analog events. Possible interpretations of isolated first analog event based on the neighboring event are listed. (B) The signature we have used to identify definite stalled forks: RUG (red-unlabeled-green) and GUR (green-unlabeled-red). Two forks moving in the same direction indicate that the fork moving in between them in the opposite direction must have stalled. (C) Ambiguous events such as RUR (red-unlabeled-red) and their possible interpretations.

A fundamental question about the regulation of fork progression in response to DNA damage is whether it is a global or local effect (Iyer and Rhind, 2013). The effect of checkpoint on origins is inherently a global response because origins distant to sites of damage are blocked from firing. However slowing of forks could be a local or a global effect. If all forks are slowed by checkpoint activation, irrespective of whether they encounter damage or not, then slowing is a global effect. On the other hand, if forks are slowed only when they encounter a lesion, then it is a local effect (Iyer and Rhind, 2013). It should be possible to distinguish between global and local regulation of fork progression by examining the effects of different lesion densities. Previously, we showed that the extent of replication slowing correlated with the density of MMS lesion, suggesting a local effect on fork progression (Willis and Rhind, 2009). However, we used a bulk replication assay, which did not allow us to directly observe replication fork rates.

Here, we have assayed checkpoint-dependent slowing of S phase in fission yeast in response to three DNA damaging drugs that activate the checkpoint at significantly different densities of lesions: methyl methanesulfonate (MMS), which mainly methylates purines and creates a relatively small adduct, 4-nitroquinoline 1-oxide (4NQO), which adds a quinoline group to purines resulting in a bulkier adduct (Galiègue-Zouitina et al., 1985; Galiègue-Zouitina et al., 1986; Sikora et al., 2010) and bleomycin, which mainly creates single strand and double strand DNA breaks (Chen and Stubbe, 2005). MMS and 4NQO create polymerase-stalling lesions and have been shown to activate the intra-S checkpoint (Larson et al., 1985; Friedberg et al., 1995; Lindsay et al., 1998; Willis and Rhind, 2009; Lopez-Mosqueda et al., 2010; Minca and Kowalski, 2011). The standard dose of 3.5 mM (0.03%) MMS causes about one lesion every 1 kb whereas a physiologically similar dose of 1 μM 4NQO causes one lesion about every 25 kb and 16.5 μM of bleomycin causes about 5 double-strand breaks per haploid yeast genome (Snyderwine and Bohr, 1992; Lundin et al., 2005; Ma et al., 2008; Asaithamby and Chen, 2009). Although these are only rough approximations of lesion density, they show that forks will encounter many more MMS lesions than 4NQO lesions or bleomycin-induced double-strand breaks. This wide disparity in lesion density allows us to address differences in global and local effects of the checkpoint. In case of 4NQO, since the lesions are rare, we expect few forks to encounter lesions. Therefore, if all forks slow in response to 4NQO then we can conclude that fork regulation by checkpoint is global in nature. On the contrary, if fork regulation is a local effect then very few forks will actually encounter the lesion and slow. Similarly since the double-strand breaks caused by bleomycin are infrequent we expect very few forks to be affected by the breaks unless the checkpoint actively regulates all forks. In case of MMS, since the lesions are frequent we expect all forks to encounter lesions, and thus to slow regardless of whether slowing is a local or global effect. By comparing the effects of these three drugs, we can differentiate between global and local effects on fork regulation by the checkpoint. We find that fork slowing is a local, checkpoint-independent effect, but that persistent fork stalling plays a more significant role in replication kinetics than previously appreciated.

## Results

### Determining the mechanism of replication slowing by DNA combing

To investigate the mechanism by which the checkpoint slows replication in response to MMS, 4NQO and bleomycin we used DNA combing. We pulsed cells in S phase with 5-chloro-2’-deoxyuridine (CldU) for 5 minutes and chased it with 5-iodo-2’-deoxyuridine (IdU) for 10 minutes, isolated and combed DNA, and visualized the CldU and IdU analogs with red and green antibodies, respectively. Figure 1 shows a sample fiber from the dataset. The fiber contains a rightward moving fork (red-green [RG]), an origin that fired during IdU labeling (green [G]), and three origins that fired during CldU labeling (green-red-green [GRG]). Further replication patterns observed in our combing dataset are shown with an interpretation of each pattern. The patterns observed were most simply interpreted as a leftward fork (green-red [GR]), a rightward fork (RG), origins that fired during CldU (GRG) or IdU (G), and terminations during IdU (red-green-red [RGR]) or CldU (red [R]). However, a more sophisticated analysis allowing the possibility of fork stalling reveals that many of these patterns—in particular termination during CldU (R)—are ambiguous, as described in the Methods section and Figure 2 and S1. For each experiment we collected about 25 Mb of DNA, which is about twice the size of the fission yeast genome, ensuring representation of most genomic loci in our analysis. From the combing data we estimated four parameters: origin firing rate, fork density, fork rate and fork stalling frequency (see Methods for details).

### Identification of stalled forks in the combing dataset using the context of neighboring events

Difficulty in identifying fork stalling events in combing data arises form the fact that stalls during the first pulse can result in ambiguous analog incorporation patterns (Técher et al., 2013). Specifically, an isolated red patch can be interpreted as an elongating fork that stalled during the first pulse or an origin with both its forks stalled or as a termination event during the first pulse (Figure 2A). Therefore, we cannot use first-pulse events alone as rigorous evidence for stalled forks. However, using double-labeled combing data we can unambiguously identify fork stalling. In particular, two neighboring forks moving in the same direction show that the fork in between them moving in the opposite direction must have stalled (Figure 2B). Thus, a red-unlabeled-green (RUG) or green-unlabeled-red (GUR) pattern is diagnostic of a fork stall. Since the fibers in our datasets average over 400 kb, we observed many neighboring forks, allowing us to robustly measure fork stalling.

Although stalled forks can be identified using the RUG and GUR patterns, some stalled forks still produce ambiguous patterns. For instance, an RUR pattern can be produced both by forks moving away from an origin and two converging forks that have stalled (Figure 2C). Thus, in order to approximate the fork stalling rate in our entire dataset, we used a probabilistic approach to estimate how many of the ambiguous patterns arise from fork stalls. We enumerated all possible incorporation patterns and classified them by the possibility that they arose from a fork stall (see Methods for details, Figure S1). We then used the stall rate from the unambiguous signals (GUR or RUG) to calculate the likelihood of ambiguous events being due to stalled forks and used that frequency for our final estimation of stall rate for each fiber.

### MMS, bleomycin and 4NQO-induced DNA damage show a similar effect on the overall replication rate

To determine the effect of MMS, bleomycin, and 4NQO—three compounds that activate the checkpoint at very different densities of lesions—on the replication rate at a population level, we analyzed cells response to them by flow cytometry. G1 synchronized cells were released into S phase with or without DNA damage, using the commonly used dose of 3.5 mM (0.03%) MMS or a dose of 1 μM 4NQO or 16.5 μM bleomycin chosen to produce a similar slowing of bulk replication (Figure S2 and S3). These doses of MMS, 4NQO and bleomycin cause lesions about once every 1 kb, 25 kb, and 3000 kb respectively (Snyderwine and Bohr, 1992; Lundin et al., 2005; Ma et al., 2008; Asaithamby and Chen, 2009). By flow cytometry, control cells completed replication by 80 minutes, while in the presence of either drug cells slowed replication, reaching only about 60% replicated by the end of the time course (Figures 3A and S4). Thus, despite the disparity in the number of lesions created by MMS, 4NQO and bleomycin, all three drugs led to a similar extent of replication slowing at the doses used. In both cases, the slowing in response to DNA damage was largely checkpoint-dependent. In the absence of the Cds1 checkpoint kinase, cells completed replication by 100 minutes even in the presence of damage, as reported previously (Figure 3B and S4) (Lindsay et al., 1998; Rhind and Russell, 1998). Although the two drugs appeared to have similar extent of slowing in wild-type, their effects in *cds1Δ* cells differed, with *cds1Δ* cells showing more checkpoint-independent slowing in MMS than in 4NQO or bleomycin (Figure 3B). We therefore examined if there is a difference in the mechanism by which DNA replication is slowed in response to MMS, 4NQO and bleomycin.

**Figure 3:**
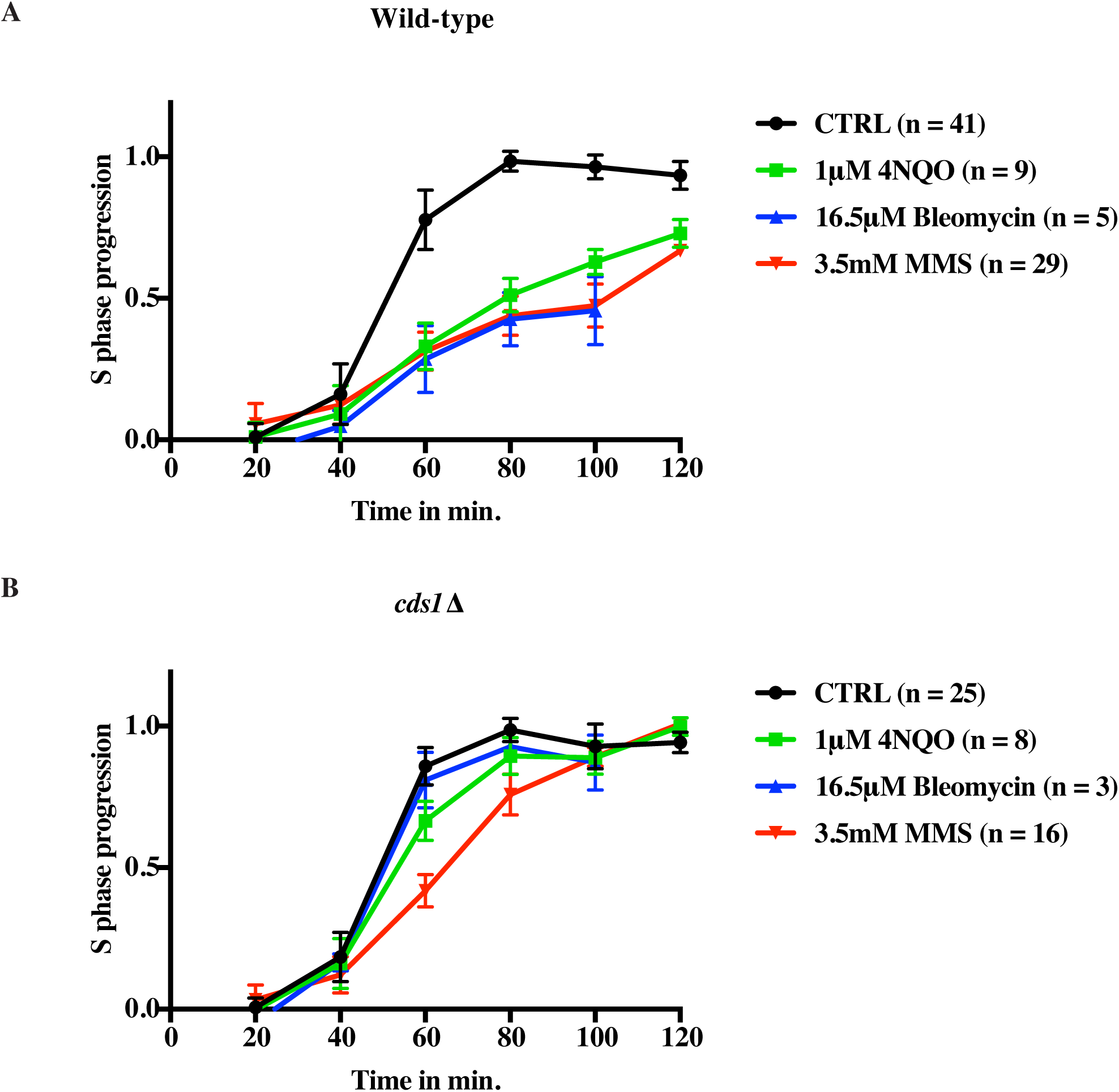
MMS-, 4NQO and bleomycin-induced DNA damage show a similar effect on the overall replication rate. (A) Wild-type cells (yFS940) show similar slowing of replication in response to 3.5 mM (0.03%) MMS, 1 μM 4NQO, and 16.5 μM bleomycin by flow cytometry. (B) Slowing in response to damage is largely checkpoint dependent by bulk assay. Wild-type (yFS940, A) and *cds1*Δ (yFS941, B) cells were synchronized in G1 and released into S phase in the presence or absence of 3.5 mM MMS, or 1μM 4NQO, or 16.5 μM bleomycin. S-phase progression was monitored by taking samples every 20 minutes for flow cytometry. The error bars represent standard deviation.

### Inhibition of origin firing is immediate in response to 4NQO and Bleomycin, but delayed in the case of MMS

As shown in Figure 3 we see similar levels of bulk slowing for all three damage treatments at 60 minutes in S phase (Figure 3 and 4A). We first analyzed the effect of DNA damage on origin firing. To measure the rate of origin firing, we directly measured the number of new origins fired during the CldU pulse (GRG patches) or IdU pulse (isolated green patches). The overall origin firing rate was calculated as the total number of origin firing events in each fiber normalized to total length of un-replicated DNA of that fiber and to the length of the analog pulse (Figure 4B). Additionally, the origin firing rate during first analog and second analog was determined separately (Figure 4D, E, F). In untreated controls, the rate of origin firing in the first analog was 2.3±0.7 origins per Mb per minute, similar to previous estimates of origin firing rates (Table S5) (Goldar et al., 2009).

**Figure 4:**
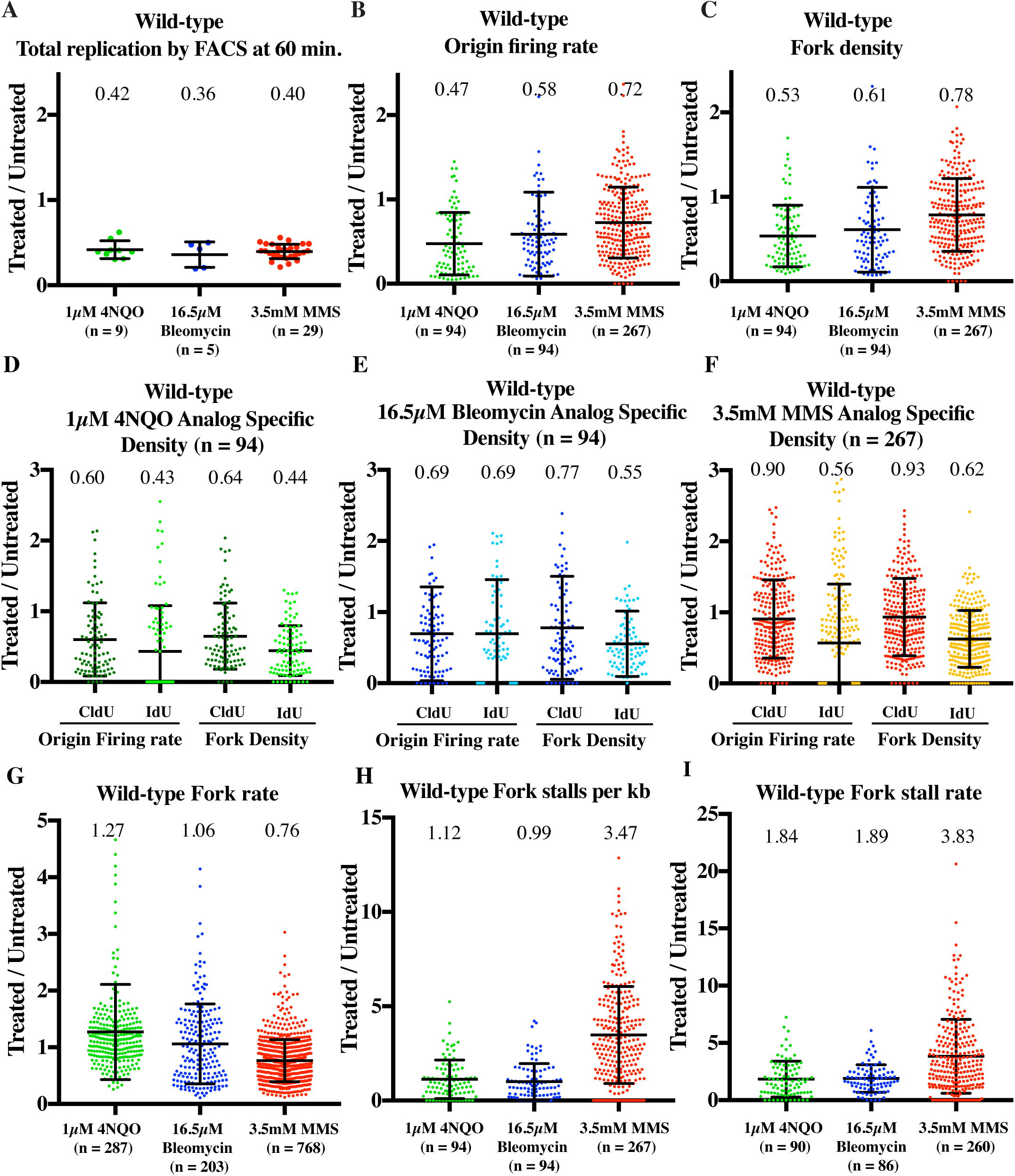
4NQO, MMS and bleomycin slow S phase using distinct mechanisms. Wild-type cells (yFS940) were synchronized in G1 and released into S phase untreated, or treated with 3.5 mM MMS, or 1μM 4NQO, or 16.5 μM bleomycin. Cells were labeled at mid-S phase with CldU followed by IdU for DNA combing. All parameters are represented as a ratio of treated v. untreated sample. (A) 4NQO, bleomycin and MMS slow S phase to similar extent by flow cytometry. Total replication by FACS was calculated at 60 minutes after release, which is close to the mid-point of analog labeling. (B) and (C) 4NQO, bleomycin and MMS show differing levels of reduction in origin firing rate and fork density. (D), (E), (F) Analog specific estimations of origin firing rate and fork density for 4NQO, bleomycin and MMS respectively. 4NQO and bleomycin treatment leads to immediate reduction in origin firing rate in the first analog while in MMS the reduction is apparent only during the second analog. (G) Fork rate decreases in response to MMS, but not in response to 4NQO or bleomycin. (H) Fork stalls per kb increases in response to MMS, but not in response to 4NQO or bleomycin (I) Fork stall rate increases in response to MMS, 4NQO and bleomycin. For calculations of origin firing rate, fork density, fork rate, fork stalls per kb and fork stall rate refer to text and methods. For each sample of each experiment, about 25Mb of DNA was collected. All error bars represent SD.

In the presence of 4NQO, the origin firing rate in wild-type cells decreased to 47% of the untreated control (p=5.17×10^−12^, t tests were used for all statistical tests) (Figure 4B and Table S5). Since, the origin firing rate only measures the origins that fired during the analog pulses, we also measured the density of active forks in the datasets in order to estimate the effect on origin firing prior to the analog pulses (see Methods for details). Fork density in 4NQO-treated cells was 64% of the untreated control (p=1.28×10^−6^) during the first analog pulse and 44% (p=3.08×10^−13^) in the second, consistent with the trend seen in origin firing rates (Figure 4D). Thus, the response to 4NQO included an immediate reduction in origin firing rate. We see a similar trend for bleomycin treated sample. The origin firing rate decreases to 58% (p=7.45×10^−5^) and the fork density decreases to 61% (p=1.33×10^−5^) (Figure 4B and 4C). Analog specific estimation shows that bleomycin treatment leads to reduction in the origin firing rate in the first analog as well as second analog to 69% (3.91×10^−4^) and shows a corresponding decrease in fork density during the first (77%, p=7.65×10^−4^) and the second analog (55%, p=8.54×10^−7^) (Figure 4E).

In case of MMS, the overall origin firing rate was reduced to 72% (p=1.18×10^−6^) (Figure 4B). Analyzing origin firing during the first and second pulse separately, we saw no statistically significant reduction in the first analog (90%, p=0.0748) followed by a stronger reduction to 56% (p=7.15×10^−11^) in the second analog (Figure 4F). Therefore, the effect of MMS on origin firing rate is delayed. This conclusion was supported by two observations. Firstly, the average fork density across both analogs in MMS showed a modest reduction to 78% (p=2.59×10^−5^) as compared to 53% (p=5.7×10^−12^) in 4NQO (Figure 4C). Second, the analog-specific fork density estimations for MMS followed a similar trend as the origin firing rate showing a greater reduction during the second analog (0.93 v. 0.62, Figure 4F). Hence, the effect of MMS-induced damage on origin firing is only manifest late in S-phase, after significant bulk slowing has already occurred, whereas 4NQO- and bleomycin-induced damage inhibit origin firing immediately in early S phase.

### Fork rate declines in response to MMS but not 4NQO or bleomycin

Since 4NQO and MMS both create polymerase-blocking lesions (Larson et al., 1985; Friedberg et al., 1995; Lopez-Mosqueda et al., 2010; Minca and Kowalski, 2011), we next studied how these drugs affect fork speed. We measured fork rate in the combing data as the length of the green track (second analog) continuing from a red track (first analog), divided by the length of the chase time (10 minutes) (Figure 1). The average fork rate in our untreated samples was 0.91±0.02 kb/minute, which is within the range of previous estimates (Conti et al., 2007; Petermann et al., 2008; Conti et al., 2010; Petermann et al., 2010; Técher et al., 2013).

In MMS, the fork rate decreases to 76% of the untreated control (p=6.13×10^−38^) (Figure 4G and S9). The reduction in fork rate was expected since the lesions are so frequent that every fork encounters about ten of lesions during the 10 minutes pulse. In case of 4NQO, the lesions are significantly less common, so only about half of the forks should encounter a lesion during the second pulse. Consistent with the low density of lesions, fork rates in 4NQO treated cells were similar to untreated cells (Figure 4G). In fact, the fork rate showed a 27% increase in the presence of 4NQO as compared to untreated cells (p=5.75×10^−4^). The dose of bleomycin used should create only 5 breaks per haploid yeast genome and therefore should not affect fork rates. Consistently we do not see any statistically significant reduction in fork rate for bleomycin treated sample as compared to untreated sample (p=0.3138) (Figure 4G).

### Fork stalling increases in response to MMS, 4NQO and bleomycin

In case of MMS treated sample, we only see a modest reduction in fork density as compared to 4NQO and bleomycin (Figure 4C). In particular, there is no statistically significant reduction in fork density in the MMS treated sample during the first analog (93%, p=0.1951) (Figure 4F) is not significant. However by bulk assay we see the same kinetics of slowing for all three damage treatments (Figure 3A and 4A). A possible explanation for this discrepancy is that forks stall during the first pulse in response to damage and thus are not observed during the second pulse (Figure 2). We therefore interrogated our combing data for evidence of fork stalling.

We first determined the absolute number of stalls per kb for each dataset (see methods for details). The absolute number of stalls per kb determines the contribution of stalled forks towards total replication slowing. In MMS treated sample we see a 3.47-fold increase in the number of stalls as compared to untreated (p=6.23×10^−27^), while we see no significant increase in stalling in response to 4NQO (1.12, p=0.604) and bleomycin (0.99, p=0.346) (Figure 4H). Thus an increase in the total number of stalls per kb in response to MMS helps explain the slowing of total replication to similar levels as compared to 4NQO and bleomycin despite having a delayed effect on origin firing rate.

Next we estimated the fork stall rate. We define the fork stall rate as the total number of stall events per fiber during the first analog pulse divided by the total number of ongoing forks in that fiber during the first analog. Although the absolute number of stalls per kb in response to 4NQO and bleomycin treatment is similar to the wild-type untreated sample, the treated samples have far fewer origins firing as compared to untreated sample (Figure 4B and 4H). Therefore the rate of stalling normalized to the origin firing rate is higher in the treated sample (Figure 4I). The average stall rate per fiber in the untreated sample was 14%. The fork stall rate showed 1.8-fold increase in response to 4NQO (p=1.2×10^−4^) and a 1.9-fold increase in response to bleomycin (p=2.26×10^−7)^, whereas MMS caused a 3.8-fold increase relative to untreated cells (p=1.55×10^−26^) (Figure 4I). Combining our stall-rate data with estimates of lesion density, we estimate that forks stall at 1.3% of 4NQO lesions and at 0.5% of MMS lesions, consistent with the fact that 4NQO creates a bulkier lesion than MMS. The stall rate per lesion is more complicated to interpret for bleomycin. Since the stalls we detect are not at the ends for our fibers, they cannot be at the sites of bleomycin-induced double-strand breaks. However, bleomycin creates about a 20-fold excess of single-strand nicks to double strand, which, and the dose we used should produce one nick about every 150 kb (Chen and Stubbe, 2005). In the absence of repair, forks would stall at 8.1% of bleomycin-induced nicks. If nicks are repaired, that rate of stalling at the remaining unrepaired nicks could be much higher. Although the stall events seem to be infrequent relative to lesion density, approximately 53% of forks in MMS-treated cells and 25% of forks in 4NQO- and bleomycin-treated cells stalled, contributing significantly to slowing, and ensuring that all treated cells had many stalled forks.

### Inhibition of origin firing is checkpoint dependent

To determine the role of the intra-S checkpoint in the observed DNA-damage dependent changes in replication kinetics, we repeated our combing experiments in checkpoint-deficient *cds1*Δ cells. In the presence of 4NQO, wild-type cells replicated 42% (p=5.55×10^−16^) as much as untreated cells, as assayed by flow cytometry at 60 minutes after release, whereas, in the absence of checkpoint, *cds1*Δ 4NQO treated cells replicated 77% as compared to untreated cells (p=5.82×10-8) (Figure 5A). 4NQO did not significantly reduce origin firing in *cds1*Δ cells (89%, p=0.206), as opposed to 47% in wild-type cells (p=5.17×10^−12^) (Figure 5B). Thus, the inhibition of origin firing in response to 4NQO is checkpoint-dependent (Figures 4B and 5B). Analog specific origin density and fork density followed similar trends as the total origin firing rate data (Figure 5D). During both the analog pulses, *cds1*Δ cells had a higher origin firing rate and fork density than wild-type cells (Figure 5D).

**Figure 5:**
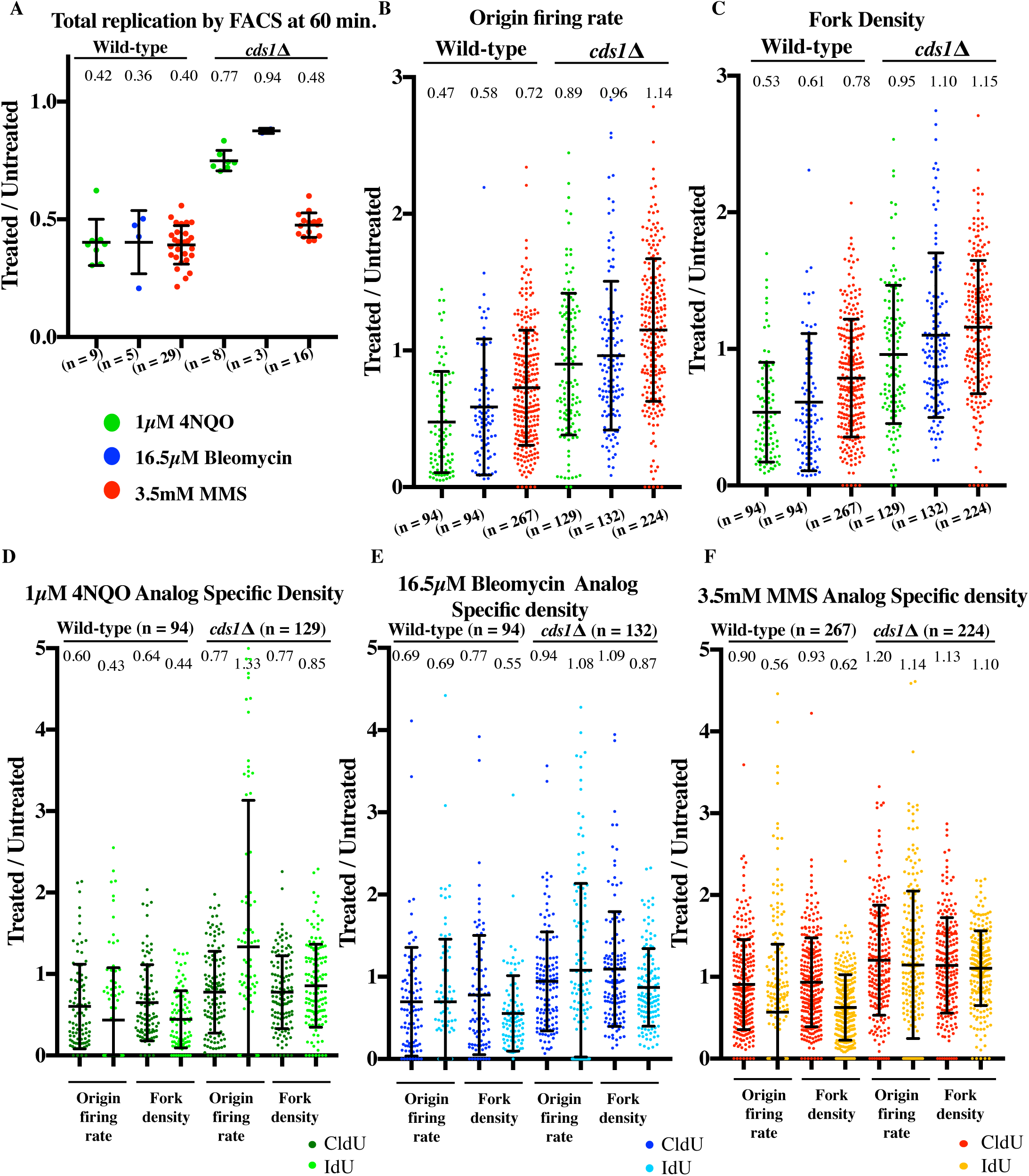
Reduction in origin firing rate is checkpoint dependent. *cds1*Δ cells (yFS941) were synchronized in G1 and released into S phase untreated, or treated with 3.5 mM MMS, or 1μM 4NQO, or 16.5 μM bleomycin. Cells were labeled at mid-S phase with CldU followed by IdU for DNA combing. All parameters from combing data of *cds1*Δ are represented as a ratio of treated v. untreated. Wild-type dataset is plotted alongside *cds1*Δ for comparison. (A) Total replication by FACS was calculated at 60 minutes after release, which is close to the mid-point of analog labeling. (B) and (C) Reduction in origin firing and fork density is checkpoint dependent for 4NQO, bleomycin and MMS. (D, E, F) Analog specific estimations of origin firing rate and fork density for 4NQO, bleomycin and MMS respectively shows higher values in *cds1*Δ than wild-type. For calculations of origin firing rate, fork density refer to text and methods. For each sample of each experiment, about 25Mb of DNA was collected. All error bars represent SD.

Likewise, inhibition of origin firing in response to bleomycin and MMS is checkpoint-dependent (Figure 5E and 5F). In wild-type cells, the overall origin firing rate was reduced to 58% by bleomycin (p=7.45×10^−5^) and 72% by MMS (p=1.18×10^−6^) treatment (Figures 4B and 5B). In contrast, *cds1*Δ cells showed no significant decrease in origin firing relative to untreated cells when treated with bleomycin (96%, p=0.389) or MMS (114%, p=0.011) (Figure 5B). Analog specific estimation of fork density and origin firing rate in *cds1*Δ treated with bleomycin or MMS showed a similar trend (Figure 5E and 5F). *cds1*Δ cells had higher fork density and origin firing rate in response to bleomycin and MMS than wild-type cells for both analog pulses (Figure 5E and 5F).

In the absence of DNA damage, the lack of Cds1 caused a 51% increase in the rate of origin firing (3.5±0.6 v. 2.3±0.6 origins firing/Mb/minute, Table S5), consistent with previous reports in other systems of checkpoint inhibition of origin firing in unperturbed S phase (Shechter et al., 2004; Petermann et al., 2010). Nonetheless, loss of Cds1 increases origin firing in damaged cells to the same level as in undamaged cells, showing that there is no checkpoint independent inhibition of origin firing (Figure S6).

### Reduction in fork rate is checkpoint independent

In MMS-treated wild-type cells, fork rate was reduced to 76% of the untreated cells (p=6.14×10^−38^) (Figures 4G and 6A). This effect was not checkpoint-dependent. In fact, at 61% (p=6.13×10^−66^), *cds1*Δ cells showed a greater reduction in fork rate than seen in wild-type cells in response to MMS (Figure 6A). Thus, the observed reduction in fork rate in response to MMS seems to be due to the physical presence of the lesions and the checkpoint activation may facilitate efficient by-pass of the lesions.

**Figure 6:**
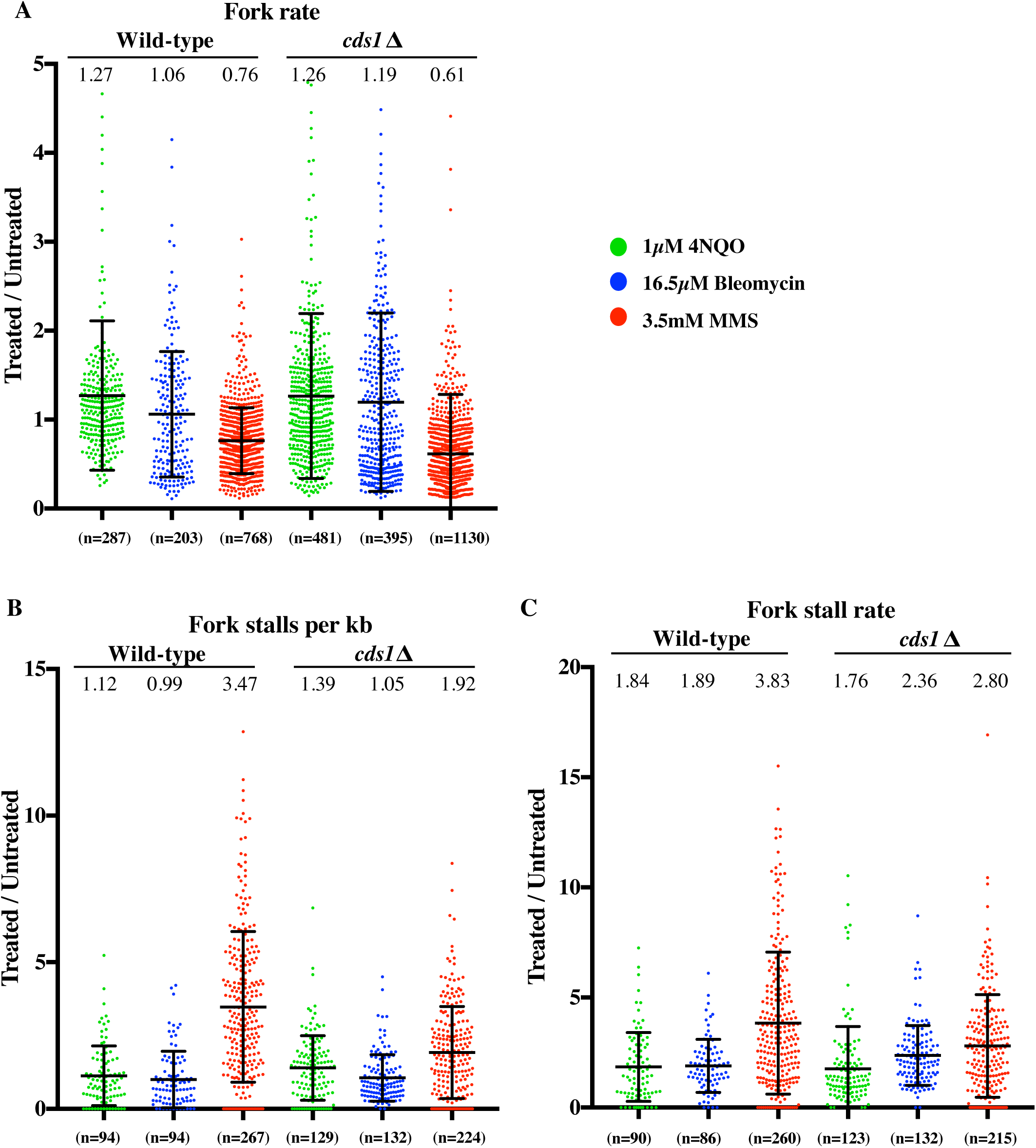
Reduction in fork rate and increase in fork stalling is checkpoint independent. (A) Fork rate, (B) fork stalls per kb and (C) fork stall rate data for *cds1*Δ (yFS941) treated with 4NQO, bleomycin and MMS. Both parameters are represented as a ratio of treated v. untreated. Wild-type dataset is plotted alongside *cds1*Δ for comparison. See methods for calculation of fork rate, fork stall per kb and fork stall rate.

Fork rates showed similar increase in *cds1*Δ. cells treated with 4NQO (126%, p=1.17×10^−5^) as they did in wild-type cells (127%, p=5.75×10^−4^) (Figures 4G and 6A). Although the lack of Cds1 does not seem to have an effect on the relative fork rate between untreated and 4NQO-treated cells, it does have a significant effect on the absolute fork rate in untreated cells. Forks moved significantly slower in untreated *cds1*Δ cells than in wild-type cells (0.72 v 0.91 kb/min, p < 10^−9^, Table S5). This difference may be an indirect effect of the higher origin firing rate and fork density in *cds1*Δ cells (Table S5). Several groups working in different systems have made the similar observation that fork rate is inversely correlated to the number of active forks, perhaps due to constrains on a limiting factor required for replication, such as the dNTP pool (Herrick and Sclavi, 2007; Herrick and Bensimon, 2008; Poli et al., 2012).

### Fork stalling in response to damage is largely checkpoint independent

Similar to wild-type cells, we see an increase in fork stalls per kb in response to MMS treatment in *cds1*Δ but not in case of 4NQO and bleomycin (Figure 4H and 6B). The fork stalls per kb shows a 1.9-fold increase in response to MMS as compared to untreated (p=4.42×10^−10^) *cds1*Δ cells. Fork stalls per kb does not change significantly between 4NQO (1.39, p=0.987) and bleomycin (1.05, p=0.839) treated *cds1*Δ cells as compared to untreated *cds1*Δ (Figure 6B).

In wild-type cells, the fork stalling rate was increased 1.8-fold by 4NQO (p=1.2×10^−4^), 1.9-fold by bleomycin (p=2.26×10^−7^) and 3.8-fold (p=1.55×10^−26^) by MMS treatment, relative to untreated cells (Figures 4H and 6C). In *cds1*Δ cells, we saw a 1.7-, 2.3- and 2.8-fold increase of the stall rate in 4NQO (p=2.59×10^−4^), bleomycin (p=7.96×10^−13^) and MMS (p=2.93×10^−15^) treated cells respectively as compared to untreated cells (Figure 6C). This increase in stall rate is slightly lower in MMS-treated *cds1*Δ cells than in wild-type cells (2.8 fold v 3.8 fold, p=9.86×10^−5^), showing that there are checkpoint-dependent and -independent contributions to stalling in response to MMS, however the difference in stall rates for *cds1*Δ treated with 4NQO (1.7 fold v 1.8 fold, p=0.746) and bleomycin (2.3 fold v 1.89 fold, p=0.01) as compared to wild-type are not as significant. Therefore the bulk of fork stalling events appears to be checkpoint-independent (Figure 6C).

### Delayed Inhibition of Origin Firing by MMS Correlates with Delayed Checkpoint Kinase Activation

To test the possibility that the later inhibition of origin firing seen in response to MMS (Figure 4F) is due to delayed checkpoint activation, and to confirm that the doses of the various damaging agents we use cause comparable checkpoint activation, we assayed activation of the Cds1 S-phase checkpoint kinase in response to MMS, 4NQO and bleomycin. Cells with an HA-tagged Cds1 were synchronized in G1 and released into DNA damage (Figure S7). Cells were harvested throughout S phase and Cds1 was immunopurified for *in vitro* kinase assays (Figure 7). We observe a significant and reproducible delay of Cds1 activation in response to MMS, relative to both 4NQO and bleomycin. These results are consistent with MMS taking longer to disrupt a sufficient number of replication forks to trigger a robust checkpoint response.

**Figure 7:**
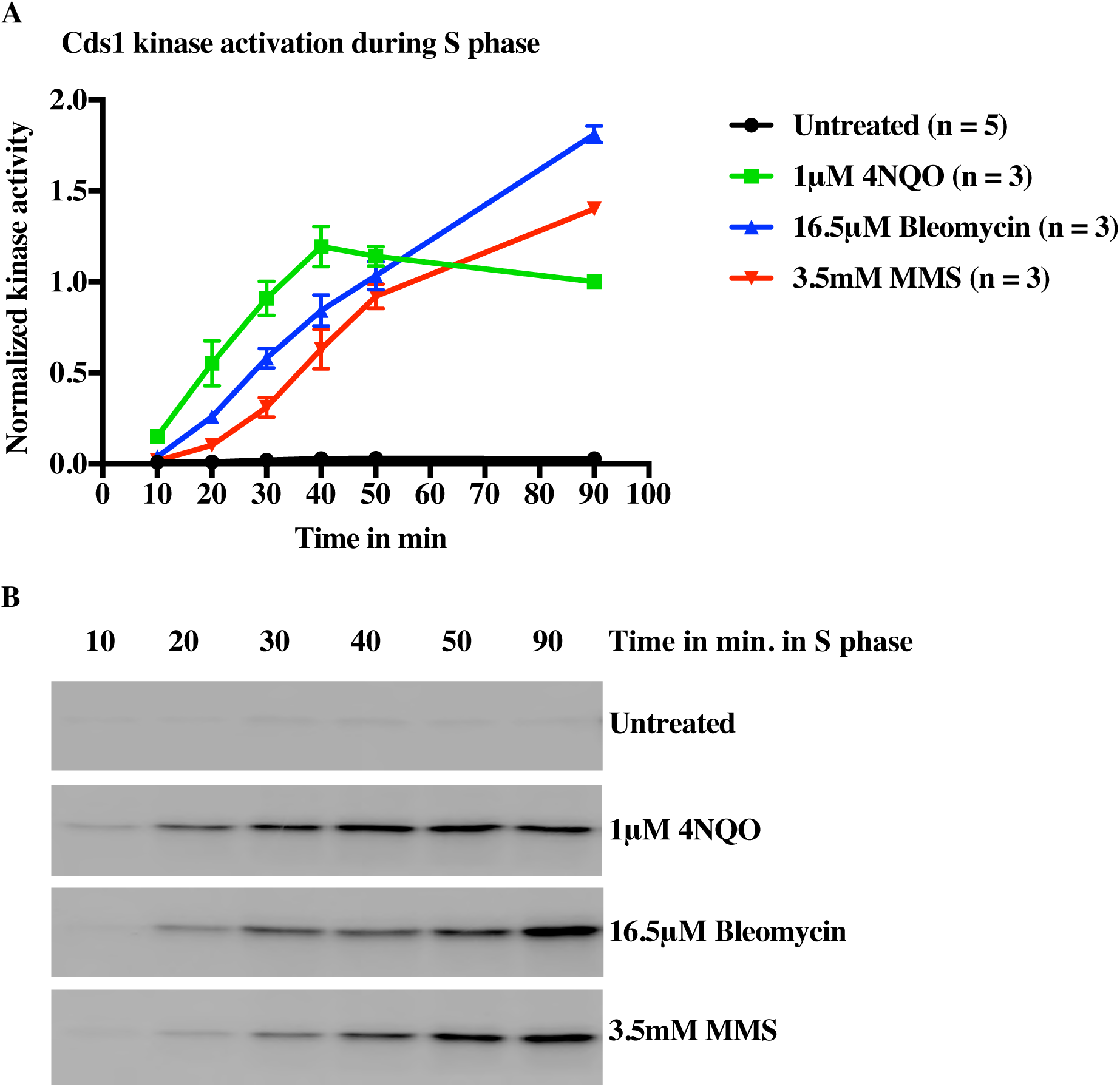
Delayed Inhibition of Origin Firing by MMS Correlates with Delayed Checkpoint Kinase Activation. A) Wild-type cells with an HA-tagged Cds1 (yFS988) were G1 synchronized, released into S phase is the presence of 1μM 4NQO, 16.5 μM bleomycin or 3.5 mM MMS, harvested at the indicated times and processed for Cds1 IP kinase assays. Average signal normalized to the 4NQO 90-minute timepoint is plotted ± standard error of the mean. B) Representative kinase assays. See Figure S8 for data from each replicate.

### Accumulation of RPA foci in response to 4NQO and MMS

To investigate the relationship between the fork stalling that we measured by DNA combing and the fate of forks in living cells, we visualized replication protein A (RPA)-GFP foci, which mark the accumulation of single-stranded DNA (ssDNA), in cells treated with MMS or 4NQO. Because RPA-coated single-stranded DNA is a trigger for the activation of the intra-S checkpoint (Zou and Elledge, 2003; Byun et al., 2005; Wu et al., 2005; Cimprich and Cortez, 2008; Xu et al., 2008), we hypothesized that stalls that accumulate substantial amounts of RPA are likely to correlate with checkpoint activation, but that stalls that lead to less RPA accumulation may be checkpoint silent.

We analyzed 100 cells for each sample and sorted them into three categories based on the intensity and number of foci: strong foci, weak foci or no foci (see Methods for details). 21% of untreated wild-type cells have some foci, consistent with previous reports (Sabatinos and Forsburg, 2015), but the majority of these are weak foci, consistent with the hypothesis that weak foci do not activate the checkpoint (Figure 8A). In the presence of damage, we saw distinctly different responses to MMS and 4NQO. In the presence of MMS, 36% of cells had strong foci (Figure 8A). Since all MMS-treated cells have stalled forks (Table S5), we conclude that only a minority of MMS-induced stalls accumulate substantial ssDNA. On the contrary, essentially all 4NQO-treated cells had strong foci, suggesting a qualitatively different nature of 4NQO-induced fork stalls, in which the majority of 4NQO-induced lesions accumulated substantial ssDNA (Figure 8A). As previously reported (Sabatinos et al., 2012), RPA did not accumulate at HU-stalled forks in checkpoint-proficient cells.

**Figure 8:**
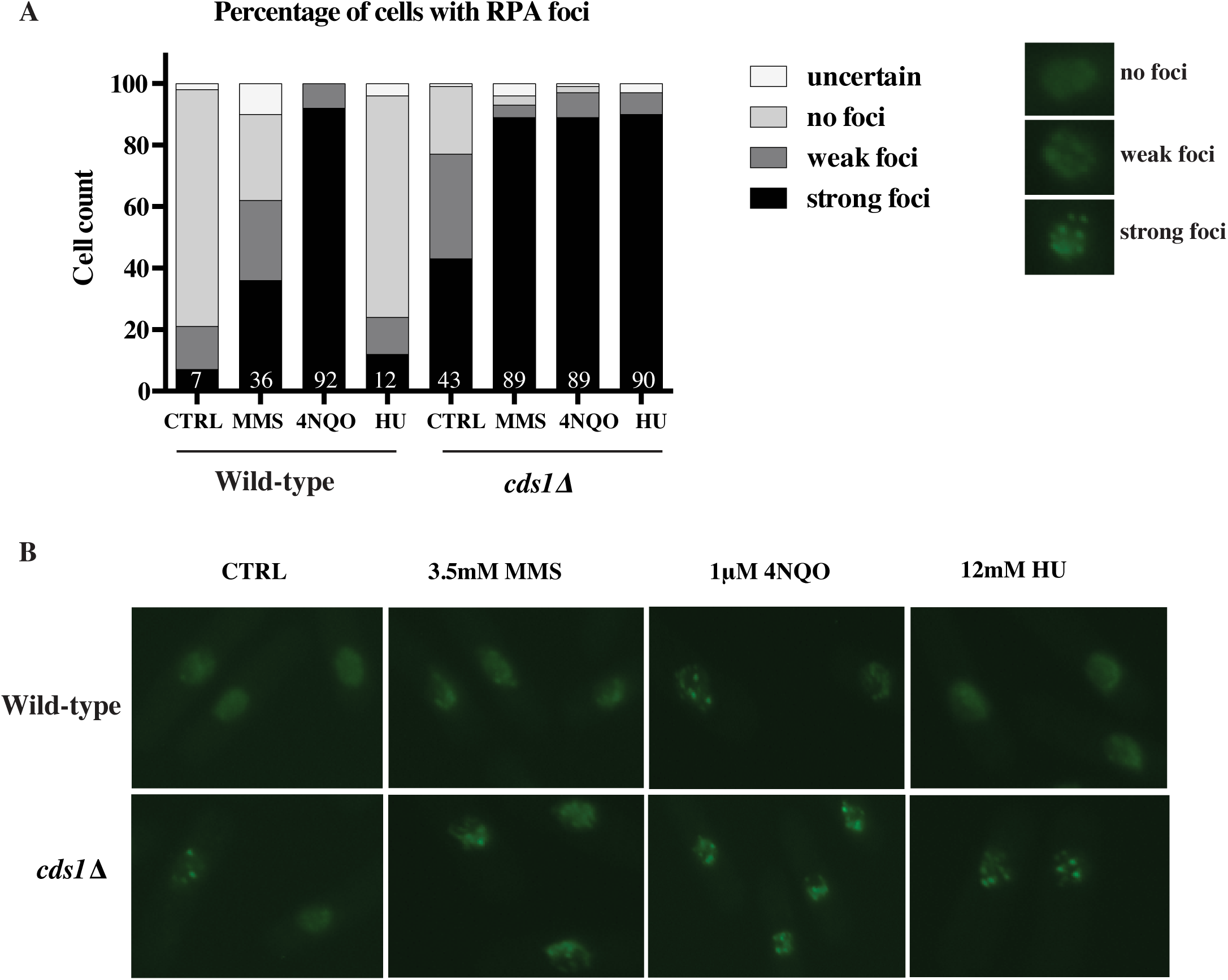
Accumulation of RPA foci in response to 4NQO and MMS. (A) Percentage of cells with RPA foci in each sample with the numbers on the bars indicating the percentage of cells with strong foci. Wild-type (yFS956) and *cds1*Δ (yFS957) cells were synchronized and released into S phase in the presence of 3.5 mM MMS,1 μM 4NQO or 12 mM HU or left untreated. Samples were collected for microscopy at 100 minutes after release. Cells were fixed and stained with DAPI and imaged in the DIC, GFP and DAPI channels. For each sample 100 cells were blinded and scored independently by two individuals. (B) Representative images for all samples quantified in (A).

In the absence of checkpoint all treated samples displayed strong foci in about 90% of cells. In particular, in MMS and HU treated cells we saw a large increase in the number of cells with strong foci in *cds1*Δ as compared to wild-type, suggesting that in response to MMS treatment the checkpoint plays a role in preventing accumulation of excess ssDNA at stalled forks, as it does in HU (Sogo et al., 2002; Sabatinos et al., 2012) (Figure 8A).

## Discussion

The regulation of DNA replication in response to DNA damage involves both inhibition of origin firing and reduction of fork speed, but relative contributions of the two effects have been unclear. Furthermore, although genetic evidence has suggested that the checkpoint regulation of fork progression is critical for genomic stability, it has been unclear if the checkpoint regulates fork speed, *per se*, and, if so, whether the regulation is at a global or local level (Wang et al., 2004; Seiler et al., 2007; Unsal-Kaçmaz et al., 2007; Szyjka et al., 2008; Iyer and Rhind, 2013). One of the reasons that these questions have persisted is that bulk methods, such as gel- or sequence-based approaches, provide an average profile of checkpoint regulation of origins and forks in response to damage and often convolve origin and fork dynamics. We have used DNA combing in fission yeast to systematically look at the effect of the checkpoint on individual origins and forks at a global scale and hence to capture the heterogeneity in response to damage. To understand the behavior of the checkpoint at a global and local scale we have used three different types of lesion inducing agents at vastly differing concentrations—3.5mM (0.03%) for MMS, 1μM for 4NQO and 16.5 μM bleomycin—which thus produce very different densities of lesions—1000 per Mb for MMS, 40 per Mb for 4NQO and 0.35 per Mb for bleomycin (Snyderwine and Bohr, 1992; Lundin et al., 2005; Ma et al., 2008; Asaithamby and Chen, 2009)—but nonetheless have very similar effects on bulk replication kinetics (Figures 3, S2 and S3). We find that the inhibition of origin firing is a checkpoint-dependent response, but one that develops more slowly in response to the many fork encounters with MMS than the less frequent but more severe fork interactions with 4NQO lesions or in response to repair of bleomycin-induced double-strand breaks. The slowing of fork progression, in contrast, is checkpoint-independent and therefore a local effect in which forks only slow when they encounter DNA damage. Finally, we discover that fork stalling, an event in which a fork stops and does not resume for the duration of the experiment, plays a significant role, with as many as 53% of forks stalling in response to DNA damage caused by MMS.

### Inhibition of origin firing is a global, checkpoint-dependent response to interactions of forks with DNA damage

Consistent with previous studies from *S. cerevisiae* (Santocanale and Diffley, 1998) and human cells (Falck et al., 2002; Sørensen et al., 2003; Merrick et al., 2004) reduction in origin firing in response to 4NQO, bleomycin and MMS is checkpoint-dependent. By flow cytometry, 4NQO, bleomycin and MMS lead to similar extent of slowing, however the combing data reveals important differences in the regulation of origin firing in response to the two drugs. In case of 4NQO reduction in origin firing is robust and immediate. The fork density during the first analog, which is a proxy for origin firing occurring prior to analog labeling, decreases to 53% (p=5.7×10^−12^) in case of 4NQO and 61% (p=1.33×10^−5^) in case of bleomycin, as compared to untreated cells (Figure 4C). Therefore, origin firing inhibition starts early in S phase in response to 4NQO bleomycin. On the contrary, in the case of MMS, fork density in the first analog is not significantly reduced (93%, p=0.074) suggesting no significant reduction in origin firing during early S phase. Furthermore, even during the second analog, the fork density is 40% higher in MMS than in 4NQO (62% v. 44%, Figure 4D and F). Therefore reduction of origin firing in MMS is delayed and modest as compared to 4NQO.

The disparity in origin firing inhibition response can be explained by the observation that a certain threshold of damage has to be met for the activation of the intra-S checkpoint. A certain number of arrested forks are necessary for checkpoint activation during S phase (Shimada et al., 2002). Therefore robust reduction in origin firing in response to 4NQO in early S phase suggests that 4NQO lesions, although less frequent than MMS lesion at the concentrations used in our studies, have more severe effects on forks and hence are more efficient at activating the checkpoint than MMS. Consistent with this interpretation, 4NQO lesions lead to a 2.6- fold higher rate of fork stalls per lesion than MMS (1.3% v. 0.5%) as detected by combing. The low density of stalls per lesion in response to MMS suggests that MMS takes longer to induce sufficient fork stalls to activate a checkpoint signal, and explains why the checkpoint mainly inhibits origin firing later in S phase. Our analysis of RPA accumulation is also consistent with this interpretation, showing that 4NQO-induced lesions accumulate RPA to a much greater extent than MMS-induced lesions (Figure 8).

### Reduction in fork rate is a local checkpoint-independent response to interactions of forks with DNA damage

The fork rate in wild-type cells is reduced in response to MMS to 76% (p=6.14×10^−38^) (Figure 4G). The fork rate is also reduced to 61% in *cds1*Δ. cells treated with MMS (p=6.13×10^−66^) (Figure 6A) consistent with previous reports (Tercero and Diffley, 2001). Thus, reduction in fork rate is checkpoint-independent and seems to be simply due to the physical presence of the lesions. In fact, by combing we see a slower fork rate in response to MMS in *cds1*Δ cells than in wild-type cells as compared to untreated (61% vs 76%, Figure 6A). The previously discussed correlation between fork rate and fork density in undamaged cells notwithstanding (Herrick and Bensimon, 2008), we do not believe this effect in MMS-treated *cds1*Δ cells is an indirect consequence of the increased origin firing, because we do not see a similar decrease in fork rate in the 4NQO-treated cells, which show much greater increase in origin firing and fork density in *cds1*Δ cells relative to wild-type cells (Figures 6A, 5D, 5F). Instead, we prefer the interpretation that the checkpoint facilitates fork progression across a damaged template. Recent work shows that even in the absence of the checkpoint, the replisome is intact at stalled forks (De Piccoli et al., 2012). Therefore, it has been speculated that the checkpoint does not affect the stability of the replisome *per se*, but instead helps maintain the replisome in a replication competent state at sites of DNA damage (Segurado and Diffley, 2008), a possibility supported by our data. In contrast, previous studies have reported similar extent of slowing in both wild-type and the checkpoint mutants in response to MMS (Tercero and Diffley, 2001; Szyjka et al., 2008). One explanation for the discrepancy could be that the previous methods—density transfer approach and BrdU-IP-seq— offer an average profile at lower resolution and mask the difference between wild-type and checkpoint-deficient cells.

Reduction of fork rate in response to MMS but not to 4NQO supports the model that slowing of forks is simply due to a transient physical slowing of forks at each lesion encountered and is not due to a global checkpoint-dependent effect on fork progression rates. Consistent with this model, we do not see a reduction in fork rate in response to bleomycin treatment (1.06, p=0.31), which activates the checkpoint robustly (Figure 4G and 7). This result is consistent with our previous observations that slowing is dependent on MMS dose (Willis and Rhind, 2009) and occurs in response to frequent UV-induced lesions, but not rare IR-induced double-strand breaks (Rhind and Russell, 1998). Since the lesions induced by 4NQO and bleomycin are 25 and 3000 times rarer than those caused by MMS, the forks are less likely to encounter damage and slow down in 4NQO-treated cells. The corollary to this conclusion is that activation of the checkpoint does not slow replication forks, as demonstrated by the fact that forks in 4NQO- and bleomycin-treated cells fail to slow despite the strong Cds1 activation and checkpoint-dependent inhibition of origin function.

### Fork stalling is a qualitatively different response to damage than fork slowing

The interaction of a fork with a DNA lesion is a first step towards recognition of damaged template during S phase and is a critical mechanism for checkpoint activation (Zou and Elledge, 2003). Activation of the intra-S checkpoint by stalled forks allows the cell to activate repair pathways, tolerate damage and prevent genomic instability (Cimprich and Cortez, 2008). Although it is believed that forks can pass polymerase-blocking lesions on the leading strand using translesion polymerases, leading-strand repriming or recombinational lesion bypass, the role of the checkpoint in such responses is unclear (Branzei and Foiani, 2005; Lee and Myung, 2008; Branzei and Foiani, 2009; Daigaku et al., 2010; Sale, 2012; Ulrich, 2012).

Here we show that forks can bypass both MMS- and 4NQO-induced lesions and that the checkpoint is not required for that bypass. However, in the case of MMS-induced lesion, the checkpoint seems to facilitate bypass, since forks move past lesions more slowly in *cds1*Δ cells as compared to wild-type (0.61% v. 0.76%, Figure 6A). We also show that, whereas forks can bypass most lesions efficiently, at a small fraction of lesions—0.5% of MMS-induced lesions, 1.3% of 4NQO-induces lesions and 8.1% of bleomycin-induced breaks—forks stall for the duration of the experiment. Given the large number of lesions throughout the genome, even these small numbers lead to a large number of stalled forks: 53% in MMS-treated cells and 25% in 4NQO- and belomycin-treated cells.

Detection of fork stalling events is complicated due to lack of consensus on how to identify them in the DNA fiber datasets (Técher et al., 2013). Generally, signal from the first (red) analog alone on a fiber (a unlabeled-red-unlabeled [URU] event) is presumed to be either an elongating fork that stalled or an origin that fired in the first pulse followed by stalling of both its forks (Figure 2A) (Merrick et al., 2004; Wilsker et al., 2008; Scorah and McGowan, 2009; Conti et al., 2010; Técher et al., 2013). Alternatively the events from the first analog alone can be interpreted as terminations (Figure 2A) (Anglana et al., 2003; Courbet et al., 2008; Letessier et al., 2011; Técher et al., 2013). However, both interpretations rely on heuristic arguments and are unable to quantitate ambiguous signals, such as URU. Therefore we developed a new, rigorous way of quantitating stalled forks in double-labeled data. We have used the context in which the first analog event occurs to define it as a stalled fork or not. As discussed in greater detail in the Results and Methods sections we have used RUG and GUR patterns as a diagnostic for fork stall event occurring during the first analog (Figure 2B). We then used a probabilistic approach to quantitate the frequency of stall events in other ambiguous patterns (Figure 2C and S1). It should be noted that we can detect a stall only if it occurs during the first (red) analog pulse and persists throughout the second (green) analog pulse.

Fork stalling does not appear to be simply an extreme example of the transient fork pausing that leads to observed fork slowing. If it were, we would expect to see a continuum of fork pause lengths form very short pauses to full stalls. Such heterogeneity would lead to a greater variation in apparent fork speeds and, in particular an increase in the asymmetry of rates in fork pairs, neither of which we see (Figure S9, S10 and Table S5). Therefore, we conclude that there are two distinct possible fate for a fork that encounters damage. It can pause briefly as it bypasses the lesion or it can stall permanently. A permanent stall does not appear to be a catastrophic event, as a majority of MMS-treated cells, which all have many stalls (Table S5), do not have strong RPA foci (Figure 8A). However, such stalls do appear to require the checkpoint to restrain ssDNA accumulation. 4NQO-induced lesions appear to cause more severe stalls, since 4NQO-treated cells have many more strong RPA foci, even though they have fewer stalls (Table S5 and Figure 8A).

The replication fork dynamics that we observe in response to MMS- and 4NQO-induced DNA damage demonstrate that forks interact with DNA damage largely in a checkpoint-independent manner. Forks are able to bypass lesions that stall the replicative polymerases with only a modest reduction in speed (Figure 6A) and are no more prone to stall at lesions in the absence of checkpoint (Figure 6B and 6C). However, stalled forks do appear to accumulate more ssDNA in the absence of the checkpoint (Figure 8). These results suggest that the major role of the checkpoint is not to regulate the interaction of replication forks with DNA damage, *per se*, but to mitigate the consequences of fork stalling when forks are unable to successfully navigate DNA damage on their own.

## Material and Methods

### General methods

The following strains used in this study were created by standard methods and grown in YES at 25°C (Forsburg and Rhind, 2006): yFS940 (*h*+ *leu-32 ura4-D18 his7-366 cdc10-M17 leu1::pFS181* (*leu1 adh1:hENT1*) *pJL218* (*his7 adh1:tk*)), *yFS941* (*h- leu-32 his7-366 cdc10-M17cds1::kanMX leu1::pFS181*(*leu1 adh1:hENT1*)*pJL218* (*his7adh1:tk*)), *yFS956* (*h*+ *leu1-32 ura4-D18 cdc10-M17 rpa1-GFP::hph-MX6*), *yFS957* (*h*+ *leu1-32 ura4-D18 cdc10-M17 cds1::ura4 rpa1-GFP::hph-MX6*), yFS988 (*h- leu1-32 ura4-D18 ade-? cdc10-M17 cds1-6his2HA*(*int*)).

### S phase progression assay by flow cytometry

Cells were synchronized in G1 phase using *cdc10-M17* temperature sensitive allele combined with centrifugal elutriation, which selects cells that have been arrested in G1 for as little time as possible (Willis and Rhind, 2011). Cells were grown to mid log phase at 25°C and arrested at 35°C for 2 hours followed by centrifugal-elutriation-based size selection at 35°C to collect cells that had most recently arrested in G1. The cells were then immediately released into S phase by shifting them to 25°C, untreated or treated with 3.5mM MMS or 1 μM 4NQO or 16.5 μM bleomycin. S-phase progression was followed by flow cytometry using a nuclei isolation protocol, as previously described (Willis and Rhind, 2011) with some minor modifications. Briefly, 0.6 O.D. of cells were pelleted every 20 minutes for 2 hours after release into S phase. Pelleted cells were fixed by resuspension in 70% ethanol and stored overnight at 4°C. Fixed cells were spheroplasted at 37°C for 1 hour in 0.6 M KCl with 1 mg/ml Lysing enzyme (Sigma # L1412) and 0.5 mg/ml Zymolyase 20T (Sunrise Science Products # N0766391). Cells were then washed with 0.1 M KCl containing 0.1% Triton X-100 followed by 20 mM Tris-HCl, 5 mM EDTA, pH 8.0. The cells were then resuspended in 20 mM Tris-HCl, 5 mM EDTA, pH 8.0 containing 250 μg/ml RNaseA and incubated at 37°C overnight. Cells were pelleted, chilled and sonicated for 7 seconds with a Sonifier (Branson Sonifier 450) equipped with a micro tip at power 5 and constant duty cycle to release nuclei. Nuclei were mixed with equal amount of 1x PBS containing 2 μM Sytox Green and analyzed by flow cytometry. S-phase progression values were obtained from the histograms as previously described (Willis and Rhind, 2011).

We have used three different drugs (3.5mM MMS, 1 μM 4NQO and 16.5 μM bleomycin) to create significantly differing lesion densities (of 1 every 1 kb, 25 kb or 3000 kb, respectively) in order to differentiate between global v. local regulation of forks. Conceivably, variable lesion density could be achieved by just titrating one of the drugs. However, we cannot achieve a 25-fold difference in lesion density (let alone a 3000-fold difference) by just titrating either drug alone. In case of 4NQO, increasing dose concentration to even 2 μM 4NQO almost inhibits replication (Figure S2), while decreasing MMS dose below 3.5mM to 0.875mM greatly reduces the effect of damage on replication kinetics (Willis and Rhind, 2009).

### DNA Combing

#### Cell labeling and plug preparation

Cells were pulse labeled with 2 μM CldU for 5 minutes and chased with 20 μM IdU for 10 minutes. For the second replicate of 4NQO dataset in wild-type and *cds1*Δ cells and all the bleomycin datasets, cells were labeled with 5 μM CldU for 5 minutes and chased with 20 μM IdU for 10 minutes. For MMS experiments were done 5 times in wild-type cells and 3 times in *cds1*Δ cells; 4NQO and bleomycin experiments were done twice in each wild-type and *cds1*Δ cells. Cells were pulse labeled at different time points across S phase for both wild-type and *cds1*Δ, however they gave similar results and hence have been combined and represented together in the figures for simplicity. Refer to Table S5 for the exact timing of analog labeling during S phase for the various replicates. Analog labeling was stopped by adding sodium azide to a final concentration of 0.1% and cooling the cells on ice. 10 O.D. of cells were pelleted for combing, frozen in liquid N_2_ and stored at −80°C. Cells were processed as previously described (Iyer et al., in press) with minor modifications. For MMS experiments the plugs were digested with proteinase K at 37°C instead of 50°C to facilitate isolation of longer fibers based on the observation that MMS creates heat-labile DNA damage (Lundin et al., 2005). Higher pH of MES (6.25 or 6.35) was used for combing instead of pH 5.4 to isolate longer fibers (Kaykov and Nurse, 2015). Combing and immuno-staining of samples were done as previously described (Iyer et al., in press). For the second replicate of 4NQO dataset in wild-type and *cds1*Δ cells, and for all the bleomycin datasets, data collection was done in collaboration with Genomic Vision, France. For experiments done in collaboration with Genomic Vision the cells were labeled and processed for combing as before, except for immuno-detection of the fibers a different set of secondary antibodies were used. The dilutions at which they were used are goat anti-rat (Abcam ab6565) 1:50, goat anti-mouse Cy3.5 (Abcam ab6946) 1:50, and goat anti-rabbit BV480 (BD Horizon 564879) 1:10.

#### Data collection

Fibers were measured in pixels using Image J (Schneider et al., 2012). Pixels were converted to kb using λ DNA as a standard. Only fibers longer than 120 kb were analyzed. For experiments done in collaboration with Genomic Vision, the fibers were manually measured using Fiber Studio software. About 25 Mb of DNA was collected for each dataset. Refer to Figure S11 and Table S5 for details of fiber lengths and dataset sizes. For each fiber, the length of each green, red and unlabeled track was manually measured. These measurements were complied into a master data table (Table S12). Analysis of this data was automated using MATLAB scripts, which are available upon request.

#### Fork rate measurement

Green tracks (containing IdU, the second analog added) continuing from red tracks (containing CldU, the first analog added) were used to determine fork rate. The length of the green track was measured from green-red (GR), red-green (RG) and green-red-green (GRG) events and divided by the length of the chase time (10 minutes). For each dataset the fork rate distribution was plotted as a histogram and was fit to a Gaussian curve. The mean fork rate was obtained from the fit (Figure S9).

We also considered that there could be a lag between the addition of analog and its incorporation into the DNA, which might lead to an underestimation of the actual fork rate. To estimate the lag we labeled the cells as previously described with 2μM CldU for 5 minutes and chased it with 20μM IdU but collected samples after different lengths of IdU incubation: 3, 4, 5, 6, 7, 8, 10, 14 and 16 minutes. However we did not find any lag in analog incorporation by plotting the lengths of IdU labeled forks across different lengths of time after IdU addition (Figure S13).

Fork rate was also analyzed by excluding forks occurring at the ends of the fiber, as broken forks may lead to an under-estimation of the actual fork rate. However, excluding the forks occurring at the ends of the fiber does not significantly affect fork rate estimations, consistent with the observation that only 6% of the forks in our dataset occur at the end of a fiber.

We note that the fork rate for wild-type untreated samples varies across different replicates (0.65±0.4 kb/min for the first three replicates v. 0.91±0.2 kb/min for the last three replicates, Table S5). We ascribe this variation to the pH of MES (6.35) used for combing the first three replicates, which we used to facilitate isolation of longer fibers. We have concluded that the stretching factor estimated from lambda DNA at 6.35 is not reliable. Hence we switched to a lower pH of 6.25 for further combing experiments. We believe that the values obtained at pH 6.25 are more reliable, since we have obtained 0.9 kb/min fork rate value from wild-type untreated samples stretched at pH 5.4, which is close to the standard pH used in most combing experiments (Figure S15) (Allemand et al., 1997; Michalet et al., 1997; Herrick and Bensimon, 1999; Bianco et al., 2012). Despite the variation in the absolute fork rate values the relative trend observed between treated and untreated sample holds true across both pH. For example consider experiments WT3-M and WT4-M, which are similar except for the pH of the combing solution used (Table S5). The absolute value of wild-type untreated fork rate differs between the experiments (0.60 kb/min vs 0.92 kb/min, Table S5). However the change in fork rate in treated v. untreated sample is 0.72 for both the experiments.

#### Origin firing rate measurement

To estimate the rate of origin firing, the total number of origin firing events in each fiber was divided by the length of the unreplicated DNA of that fiber and by the total length of the analog pulses (15 minutes). The total number of origins firing in the fiber is the sum of origins that fire during the first and the second analog. Origins that fire during the first analog are identified as GRG events. Origins that fire during the second analog are identified as isolated green events. However, origins that fire in the first analog will appear as GRG only if both its forks progress into the second analog labeling. In the event of a unidirectional fork stall during the first analog pulse, origins that fire will appear as GR or RG. An origin may also appear as an isolated red event if both its forks stall during the first analog pulse. Therefore, the origin firing rates were corrected by accounting for the probability of forks stalling during the first analog pulse which would have disrupted GRG events, based on the measured fork stall rate (see below for stall rate calculation).

#### Fork density measurement

Origin firing rate captures the origins that fire during the course of the analog pulses. However, the rate of origin firing prior to labeling influences the density of forks during the pulses and provides a parallel measure of origin activity. Therefore, we measured fork density to assess origin firing rate during the period prior to our S-phase labeling pulses. Fork density, the total number of forks in each fiber was divided by the length of the unreplicated DNA of that fiber, and was calculated independently for the two labeling pulses. The basic strategy for measuring the fork density during either analog pulse is straight-forward. For the first pulse, it is done by counting the following events in each fiber and dividing by the length of the unreplicated DNA of that fiber: the number of origins firing (GRG) and terminations (R and RGR) during the first pulse multiplied by two, since each origin or termination comprises two forks, and the unidirectional forks (GR and RG) counted once. Likewise, for the second analog pulse it is done by counting the following events in each fiber and dividing by the length of the unreplicated DNA of that fiber: the number of origin firing during the second pulse (G), forks from origins firing in the first pulse (GRG) and terminations during the second pulse (RGR) multiplied by two, and the unidirectional forks (GR and RG) counted once. We also calculated the total fork density for each fiber, which is done by counting following events in each fiber and dividing by the length of the unreplicated DNA of that fiber: number of origins firing during each analog (GRG and G) and terminations (R and RGR) multiplied by two, and the unidirectional forks (GR and RG) counted once. However, these calculations are confounded by forks that stall during the first pulse, leading to the misclassification of events (Figure 2). To account for such stalling events, we took a probabilistic approach, in which the measured stall rate is used to more accurately estimate fork density, as described below.

#### Fork stall rate measurement

We can estimate the fork stall rate only during the first analog pulse. Stalls occurring during the second analog pulse simply reduce the length of the second analog incorporation track and thus get interpreted as a reduction in fork rate. Fork stall events occurring during the first analog pulse can lead to ambiguous incorporation track patterns (Figure 2A). For instance, an isolated first analog event is open to any of the following three interpretations: an unidirectional elongating fork that stalled, or an origin firing event for which both forks stalled, or a termination event. However, two apparently unidirectional forks moving in the same direction on a fiber must have had the fork in between them stall (Figure 2B, Figure S1A). Such unambiguous stall events leave signature red-unlabeled-green (RUG) or green-unlabeled-red (GUR) patterns. On the other hand the signature for two unstalled forks is green-unlabeled-green (GUG) (Figure S1A). To estimate the apparent stall rate in the dataset we counted every RUG and GUR event and divided it by the sum of GUR, RUG and GUG events (Figure S1A). The GUR and RUG events are included in the denominator because every stall event represents a loss of one fork. For example, consider an origin (GRG) on a fiber. If one of its fork is stalled then it will appear as GR or RG and we would calculate the stall rate as 50% since one fork is stalled but ideally there should be two (one fork which we can visualize and one which is stalled).

Although the unambiguous events allow us to determine the stall rate with certainty, not every isolated first analog event is flanked by second analog events to help us determine whether it is stalled or not. For example consider Figure 2C, which shows two forks moving away from one another. It can be interpreted as forks moving away from a single origin or as two origins with stalled forks on their inner side. To estimate the stall rate across the ambiguous events we used a probabilistic approach. We considered all combinations in which ambiguous events could occur (e.g. UR, RU, RUR, GRU, URG, GRUR, RURG, URUR, RURU, RURUR etc.) (Figure S1B). The ambiguous events were assigned a probability of being either stalled or not, using the apparent stall rate as the probability of a fork stalling (Figure S1B). For example, consider red-unlabeled-red (RUR) event represented in Figure 2C, if the leftward moving fork is interpreted as an origin with stalled rightward fork then it automatically implies that the rightward moving fork on the fiber is also an origin with stalled leftward moving fork. Therefore the probability of both the events being considered as origins with a stalled fork, is the product of two stall events occurring at the same time i.e. square of the apparent stall rate. If the apparent stall rate is 10%, then RUR events are assigned a 1% probability of as having two stalled forks. The probabilistic interpretation of the ambiguous events also changes the absolute number of forks in the dataset. For example according to interpretation 1 in Figure 2C there is only one origin with two elongating forks on the fiber. However according to interpretation 2 there are two origins with a total of 4 expected and 2 apparent forks. To estimate the stalls per kb for each fiber the unambiguous stall events (RUG and GUR) in each fiber were combined with the fraction of the ambiguous events that were predicted to be stalled and this total was divided by the length of the un-replicated DNA. To estimate the stall rate per fiber the unambiguous stall events (RUG and GUR) in each fiber were combined with the fraction of the ambiguous events that were predicted to be stalled and this total was divided by the total number of ongoing forks in the first analog, which was calculated as the sum of GR, RG, and R events (counted once or twice based on whether they were interpreted as elongating forks or as an origin with a stalled fork) plus two forks for each GRG.

The fork stall rate in untreated wild-type and *cds1*Δ cells is about 15% (Table S5). This result reveals an unexpectedly high rate of spontaneous fork stalling. To exclude the possibility that the high stall rate is a result of previously identified combing artifacts (Demczuk and Norio, 2009), we re-analyzed our dataset, excluding the labeled events occurring at the ends of the fibers, where such artifacts can occur. However the stall rate estimations remain unchanged after re-analysis (Figure S14). We cannot rule out the possibility that the observed fork stall rate is increased by the thymidine analogs used to label the DNA. Nonetheless, since we normalize our quantitation of fork-stalling to an untreated sample, our results are internally controlled. Furthermore, double-labeled combing studies done in mammalian cells show similar rate of fork stalling in the untreated sample (Merav Socolovsky, personal communication).

#### The effect of analog addition on replication kinetics

Addition of analog early in S phase causes slowing of bulk replication, probably due to checkpoint-dependent origin inhibition (Figure S16A). Therefore, we have labeled cells in a manner so as to incur minimal effect on S phase progression due to analog addition (Figure S16B). By adding analogs at different time points across S phase and examining the effect on S phase progression by FACS, we chose to add analog at 50 minutes after release and harvest cells for combing by 65 minutes after release for wild-type (Figure S16B). Furthermore, all of our interpretations are based on comparison to analog-treated cells, so any analog-specific effects are controlled for.

#### Cds1 kinase assay

To estimate the Cds1 kinase activity S phase progression assay was done in triplicate as described earlier in response to 3.5 mM MMS, 1 μM 4NQO, and 16.5 μM bleomycin in yFS988 strain (Figure S7). Approximately 10 OD of cells were pelleted at 10, 20, 30, 40, 50, and 90 minutes in S phase for measuring Cds1 activity. Cells were lysed by bead beating in 400μl ice-cold lysis buffer (150 mM NaCl, 50 mM Tris pH 8.0, 5 mM EDTA pH 8.0, 10% Glycerol, 1%IGEPAL CA630, 50 mM NaF, freshly added 1 mM Na_3_VO_4_ and protease inhibitor cocktail (Sigma 11836170001)) for 15 minutes at 4°C. Lysate was cleared by centrifuging at 1000g, 5 minutes and combined with anti-HA Ab conjugated agarose beads (Pierce 26181) and incubated with constant mixing at 4°C for 4 hours. Beads were washed twice with lysis buffer and twice with kinase buffer (5 mM HEPES pH 7.5, 37.5 mM KCl, 2.5 mM MgCl2, 1 mM DTT). The sample was then split into two portions, one was used to estimate the amount of Cds1 pulled down by western blot and the other was processed to estimate the kinase activity. For western, the Cds1 was eluted off the beads by boiling in 2x SDS PAGE gel loading dye. Cds1 was detected by using rabbit anti-Cds1 Ab at 1:1000 and anti-rabbit HRP conjugated secondary Ab at 1:10000. For kinase assay, the beads were re-suspended in 10 μl 2x kinase buffer, 0.5 μl 10 μCi/μl γΡ32- ATP, 2 μl 1 mM ATP, 5 μl 1 mg/ml myelin basic protein and incubated at 30°C for 15 minutes. The reaction was quenched by adding SDS PAGE gel loading buffer and boiling at 95°C for 5 minutes. Kinase reactions were run on a 15% gel normalized to the amount of Cds1 pulled down in each lane. The gel was dried under vacuum and exposed to phosphoimager screen for 48 hours. The screen was scanned on Typhoon FLA-9000 and quantitated using ImageJ (Schneider et al., 2012). Asynchronous culture of yFS988 treated with 3.5 mM MMS for 4 hours was used as a control to normalize across gels and blots.

#### RPA foci estimation

In order to estimate RPA foci formation due to damage, S-phase progression assays were performed as described above with strains expressing GFP-tagged Rpa1 (yFS956 and yFS957). Cells were collected at 100 minutes after release from untreated, 3.5 mM MMS, 1 μM 4NQO and 12 mM HU treated cultures and fixed in ice-cold 100% methanol and stored at −20°C for at least 20 minutes. Fixed cells were washed thrice with 1x PBS and re-suspended in 10 μl Vectashield mounting medium with DAPI at 2 μg/ml. Cells were visualized using a Zeiss Axioskop 2 Plus epifluorescence microscope with 100X Plan-NEOFLUAR oil objective and imaged using SPOT monochrome cooled-CCD camera. Images were analyzed using ImageJ and Microsoft Excel. Image of each nuclei in the GFP channel was cut and pasted into cells in an excel file. The nuclei were then blinded, randomly sorted and scored independently by two individuals. 100 nuclei were analyzed for each sample. The nuclei were scored into three categories: uncertain, foci negative, and foci positive. Foci positive cells were further characterized into nuclei with few weak foci (1 or 2) or containing many weak foci, strong foci or a combination of strong and weak foci.

## Acknowledgments

We thank Atanas Kaykov for suggestion regarding the preparation of silanized coverslips and the pH of the combing buffer, which allowed us isolate longer fibers, and Merav Socolovsky and Shankar Das for their helpful suggestions regarding the combing protocol. We also thank members of the Rhind lab for constructive comments of the project and this manuscript. This work was supported NIH grant GM098815 to NR.

---

**Figure S1:**
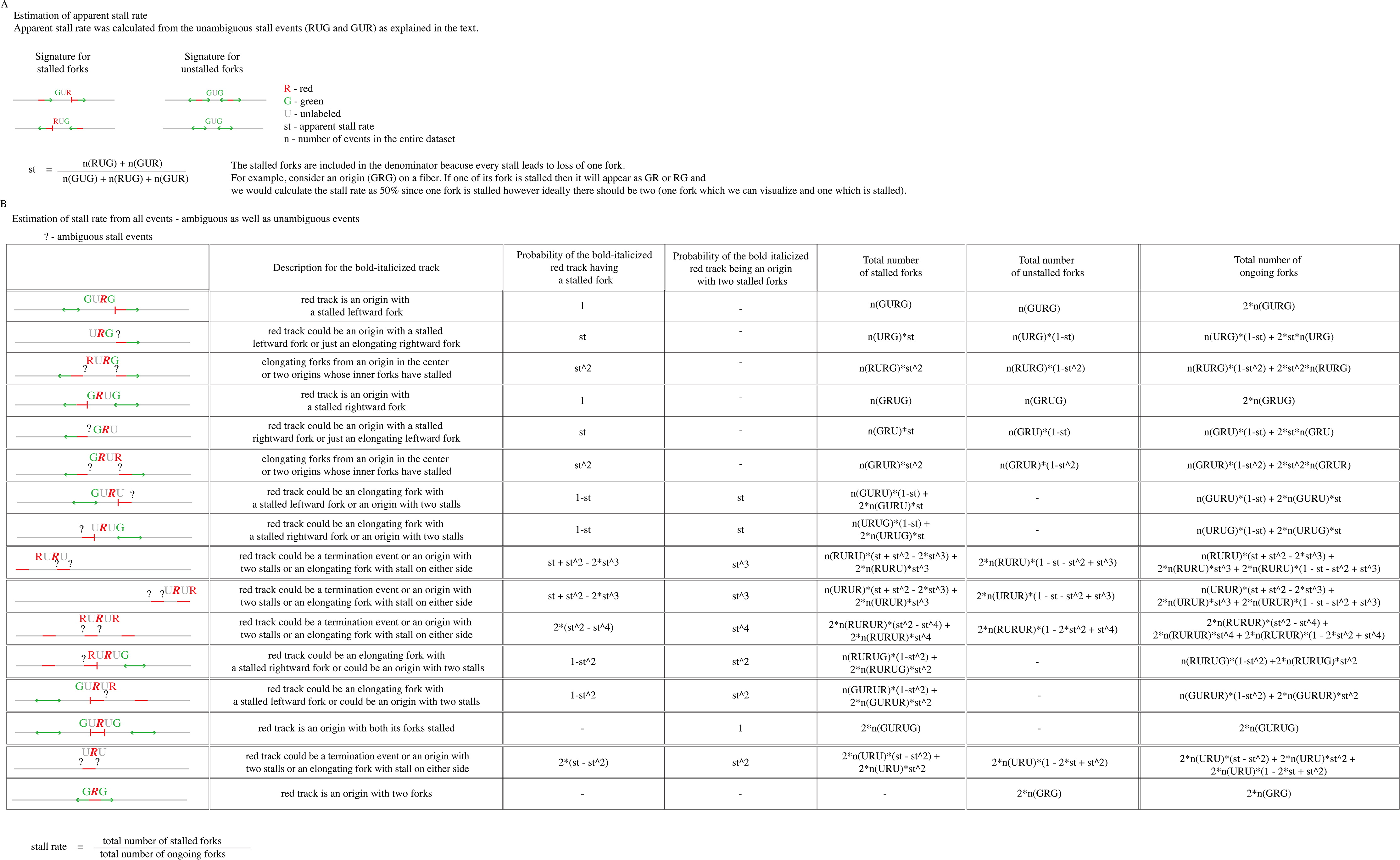
Schematic explanation of stall rate estimation.

**Figure S2:**
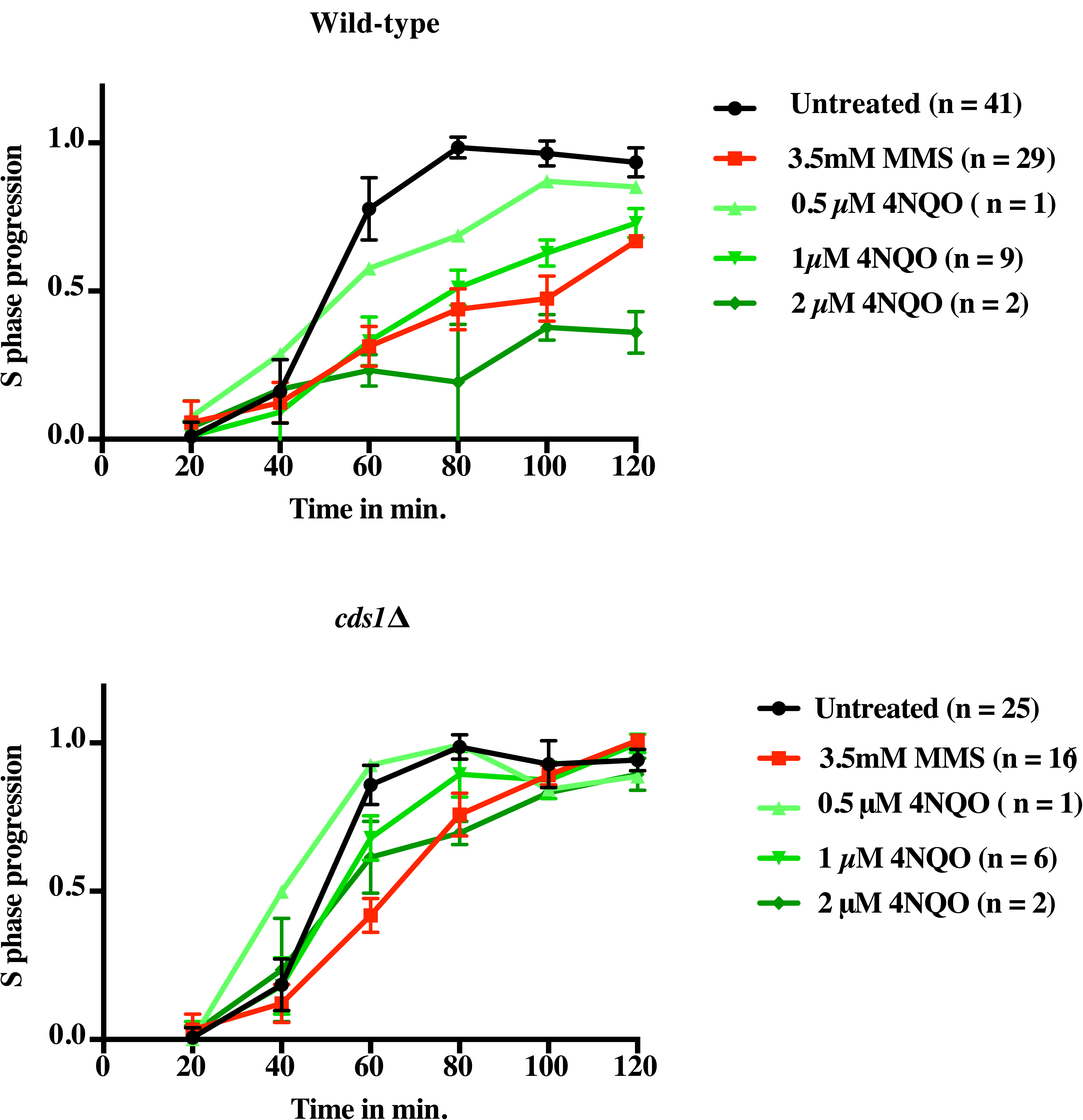
Titration of 4NQO. Wild-type (yFS940) and *cds1*Δ (yFS941) cells were synchronized and released into S phase with 3.5 mM MMS, 0.5 μM, 1 μM, 2 μM 4NQO or left untreated. S-phase progression was monitored by taking samples every 20 minutes for flow cytometry.

**Figure S3:**
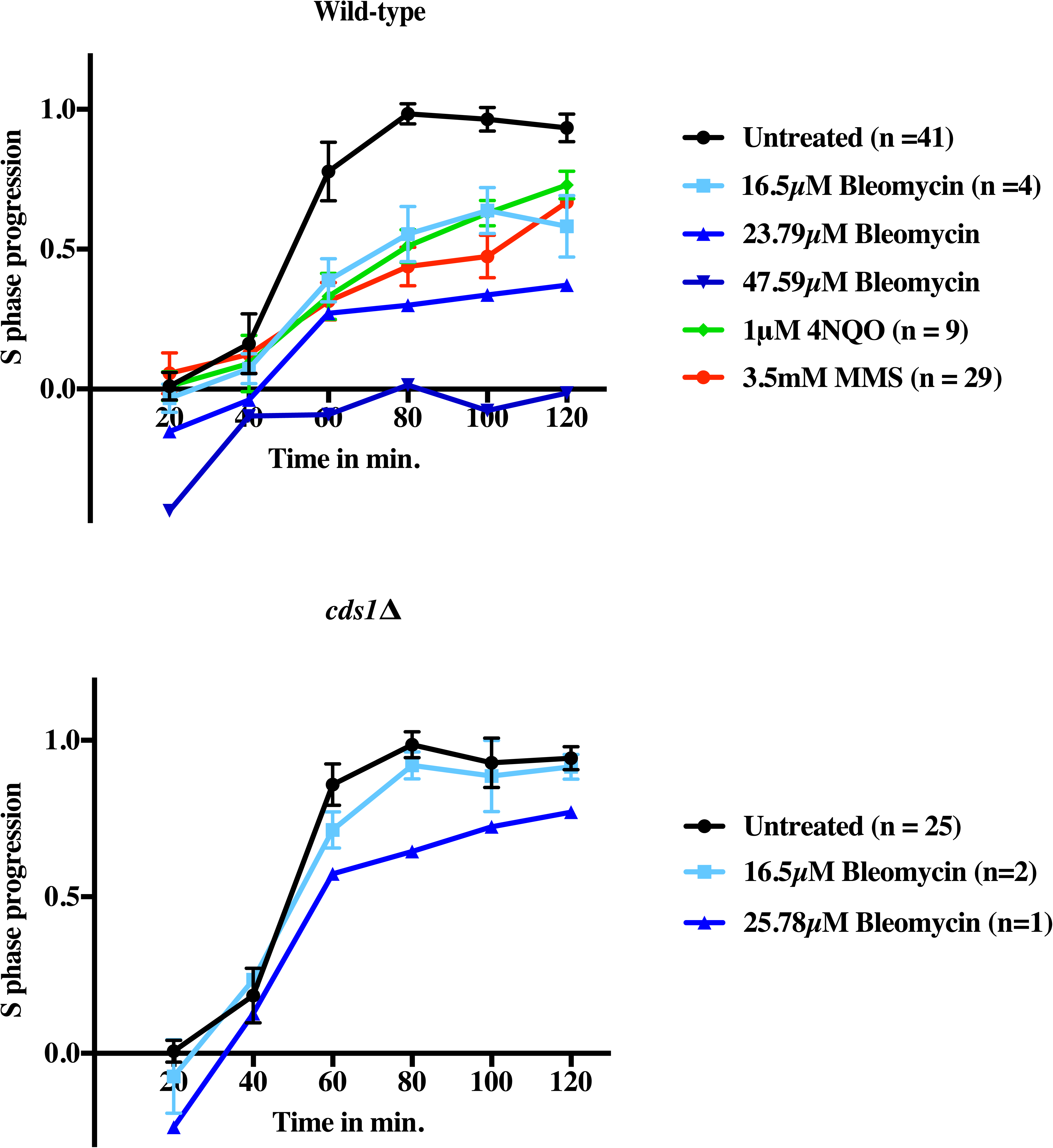
Titration of Bleomycin. Wild-type (yFS940) and *cds1*Δ (yFS941) cells were synchronized and released into S phase with different concentrations of bleomycin 16.5 μM, 23.79 μM, 47.9 μM or left untreated. S phase progression was monitored by taking samples every 20 minutes for flow cytometry.

**Figure S4:**
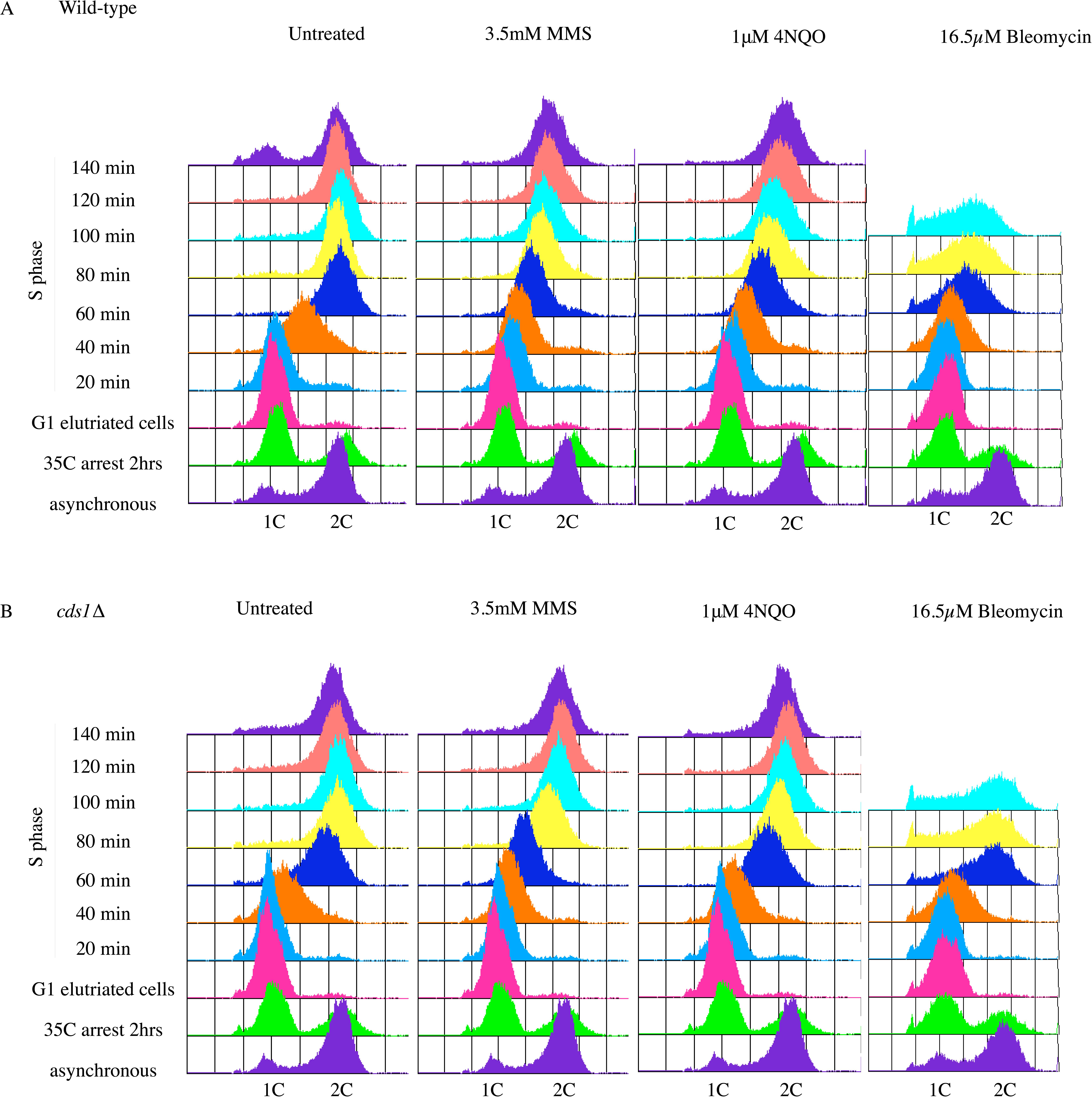
S phase progression by FACS. WT (yFS940) (A) and *cds1*Δ (yFS941) (B) cells were synchronized in G1 phase using cdc10-M17 temperature sensitive allele followed by elutriation. Elutriated G1 cells were released into permissive temperature untreated or treated with 3.5 mM MMS or 1 μM 4NQO or 16.5 μM Bleomycin.

Table S5: Summary of dataset information and results.

**Figure S6:**
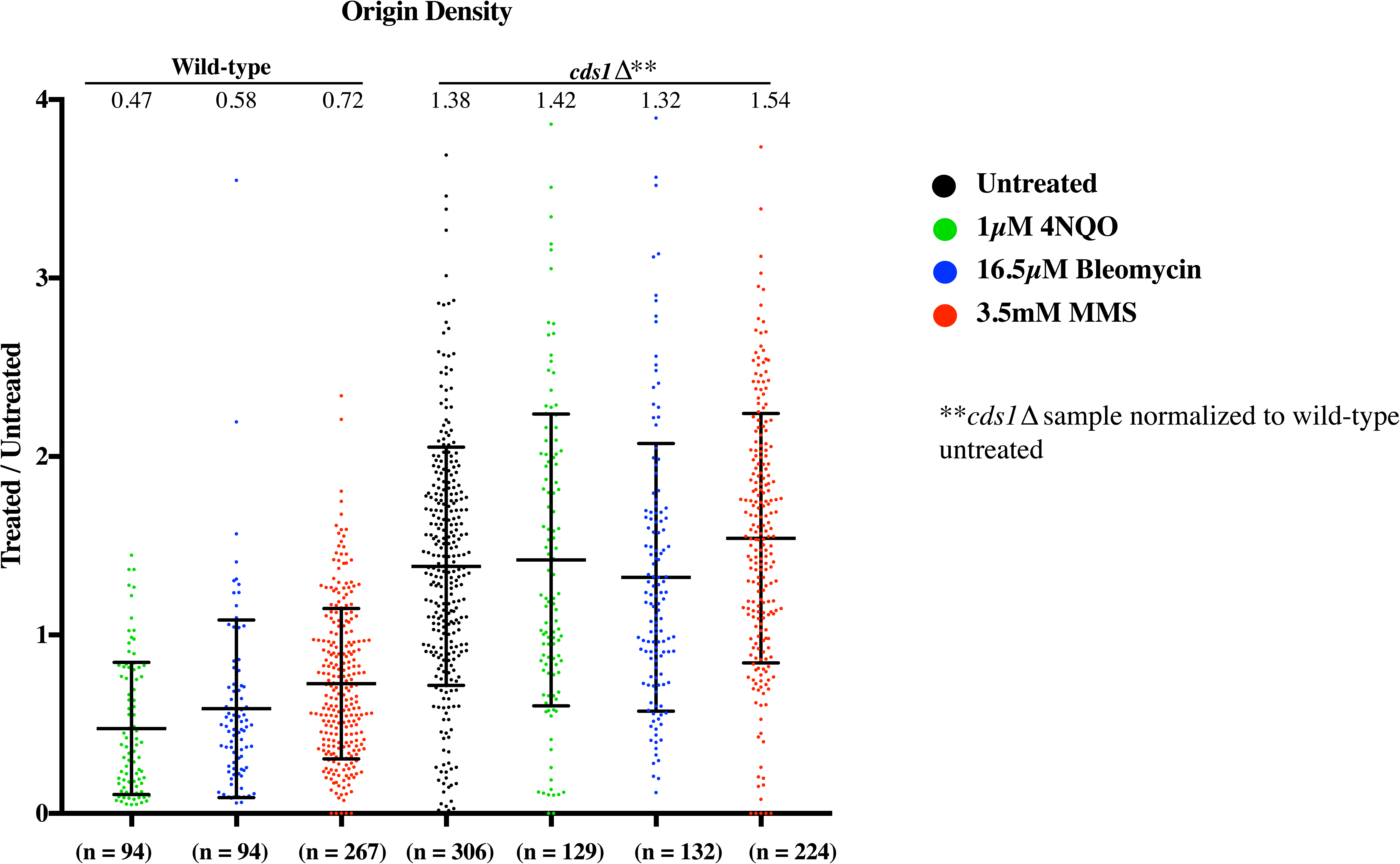
Origin density from *cds1*Δ sample normalized to wild-type untreated sample.

**Figure S7:**
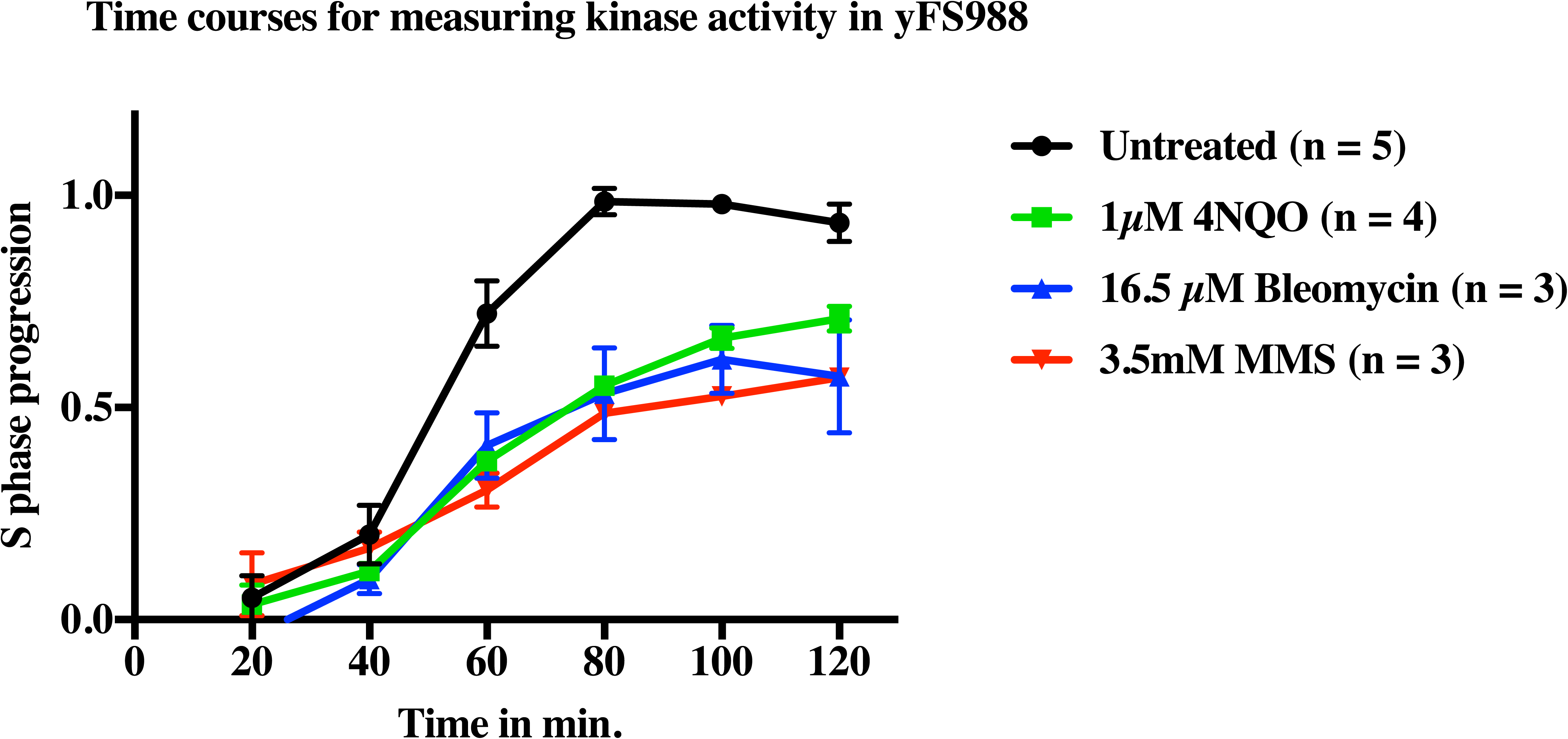
S phase progression by FACS of time courses used for kinase activity measurement.

**Figure S8:**
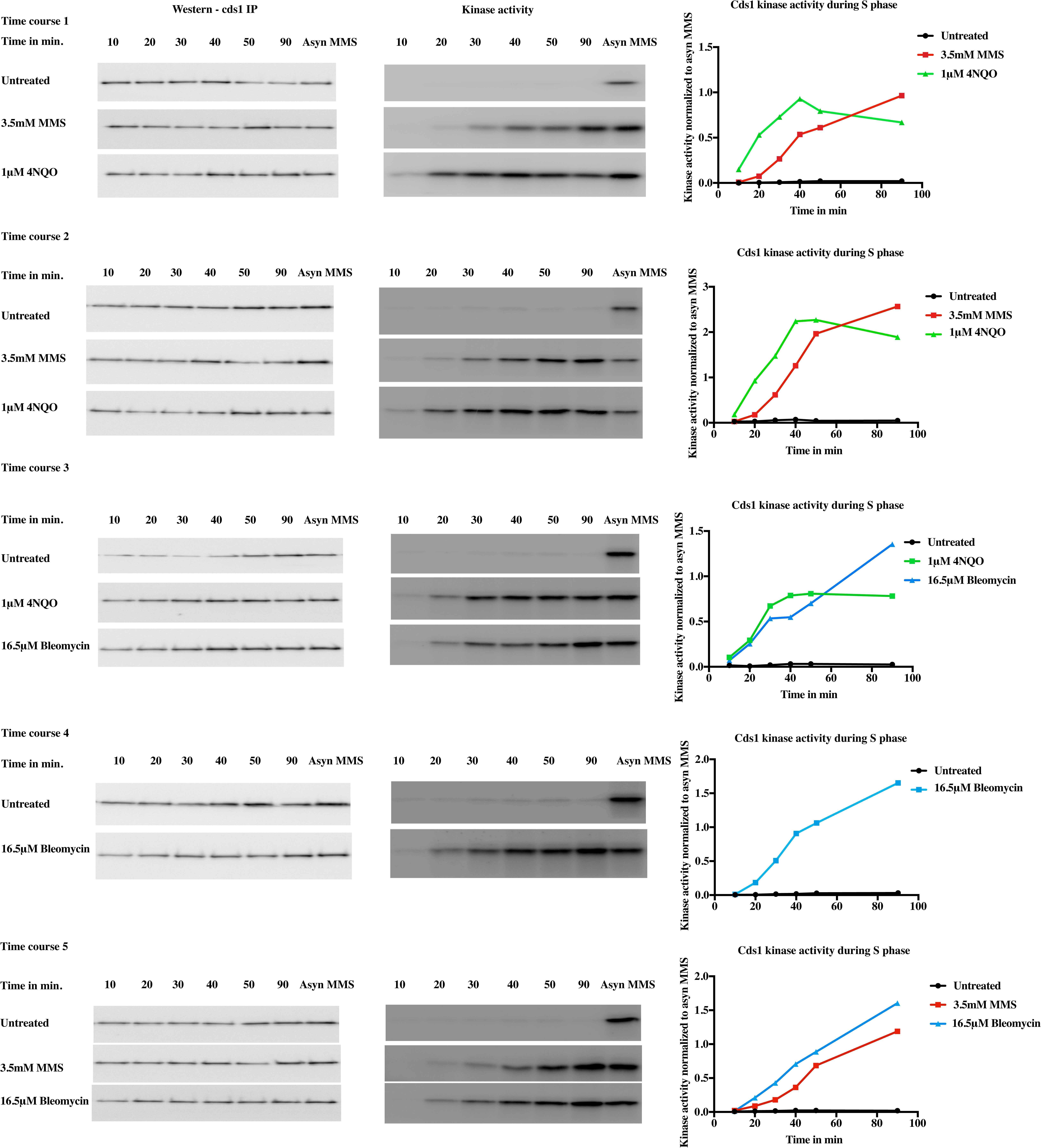
Kinase assay to measure Cds1 activation in 5 time courses. For each time course, Cds1 western was done to estimate the amount of Cds1 pulled down in each condition. Kinase assay reactions were run normalized to the amount of Cds1 pull down. Signal from each lane of the kinase assay gel was normalized to the asynchronous MMS sample to get the normalized kinase activity

**Figure S9:**
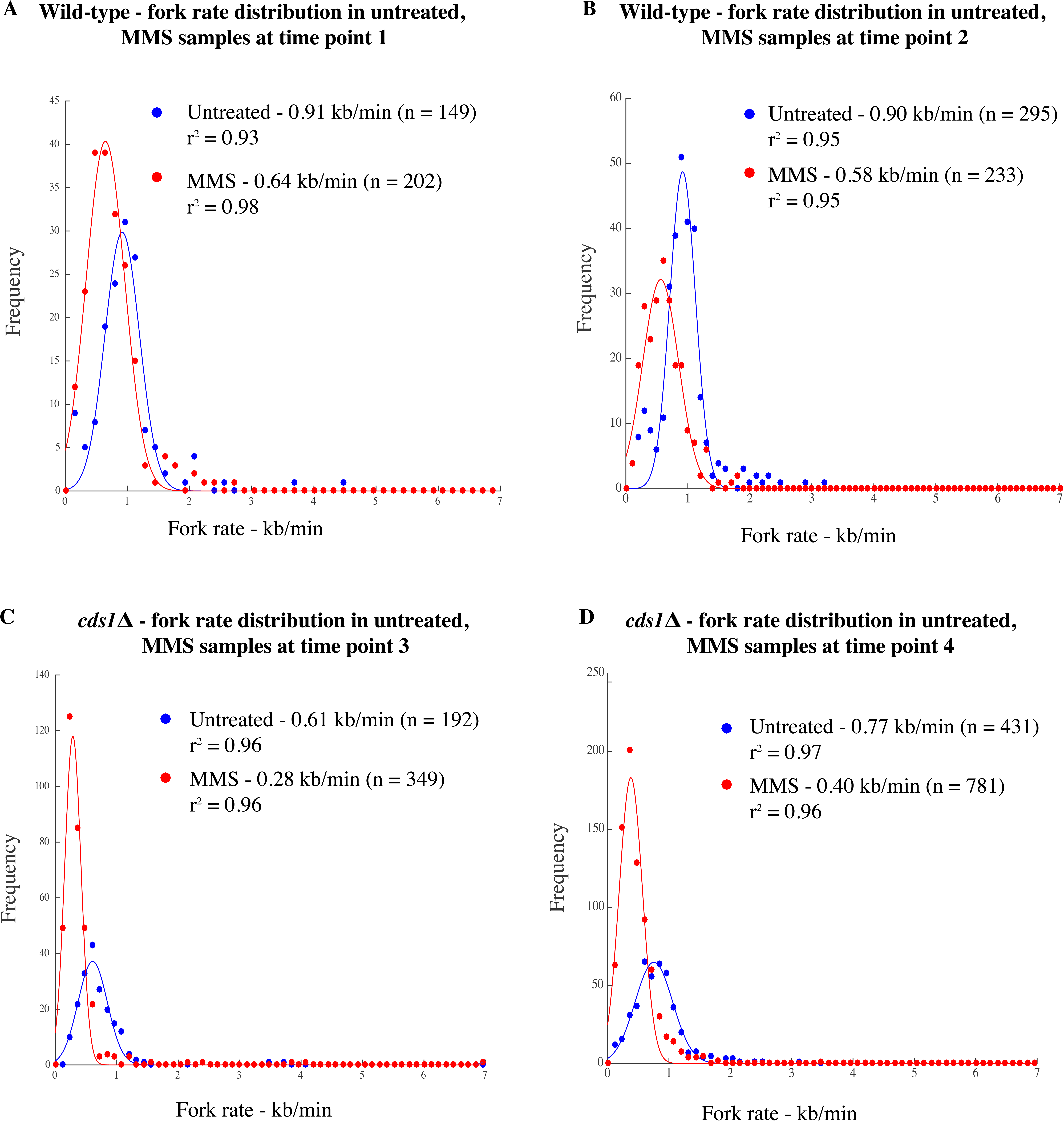
Fork rate distribution. Fork rate distribution in wild-type and *cds1*Δ cells untreated or treated with MMS from a single experiment at time points 1 and 3, respectively (A and C) and from two experiments at time points 2 and 4, respectively (B and D). For the exact timings refer to Table S5. Each distribution of fork rate was fit to a Gaussian curve. Mean fork rate was obtained from the fit.

**Figure S10:**
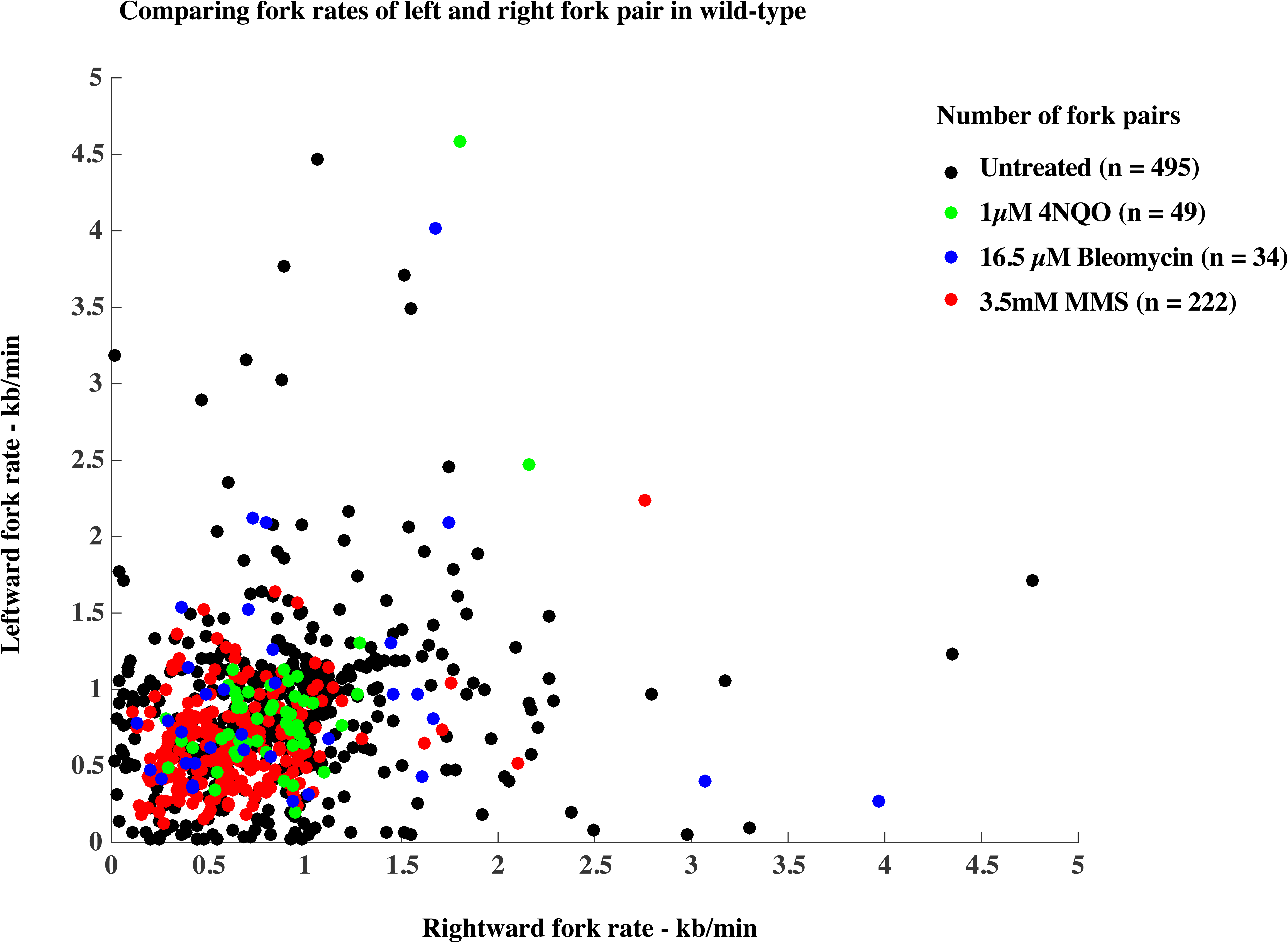
Comparing fork rates of left and right fork pairs in wild-type (yFS940) does not show increased asymmetry in treated samples as compared to untreated sample.

**Figure S11:**
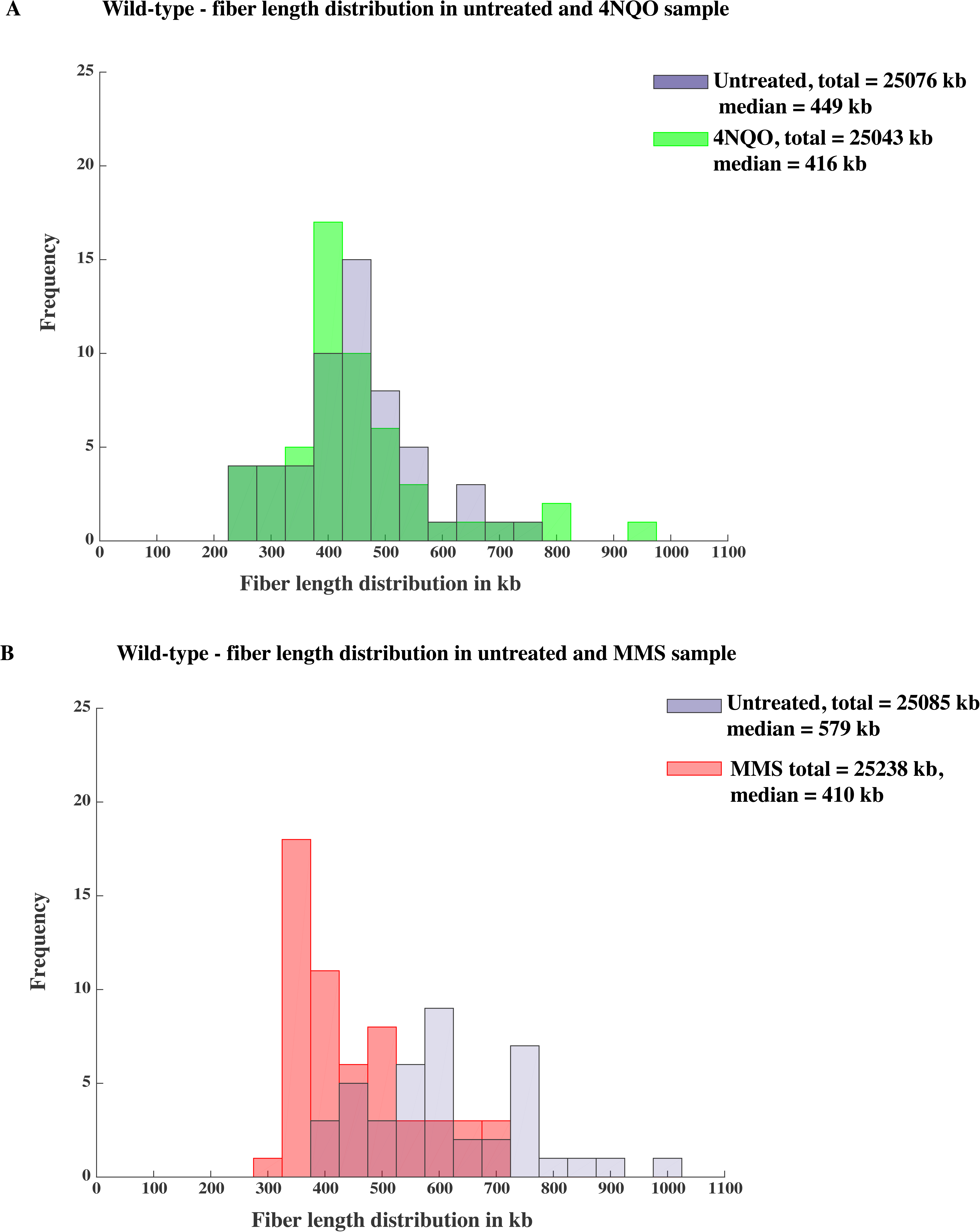
Fiber length distribution. Length of fibers from wild-type 4NQO (A) and MMS (B) samples from a single experiment. Approximately 25 Mb of total DNA was collected for each sample (Table S5).

Table S12: Raw data files from all datasets containing measurements of each fiber.

**Figure S13:**
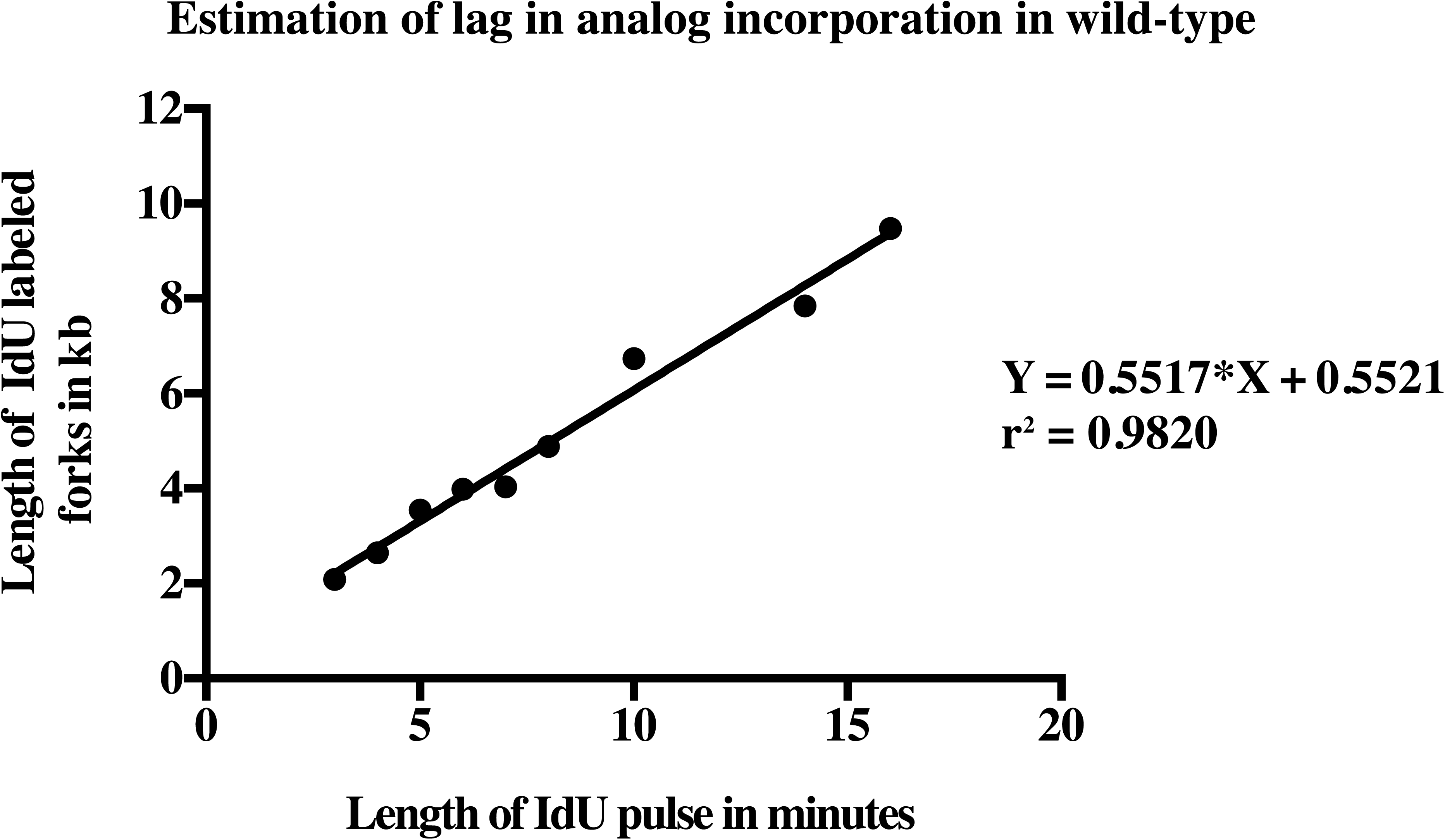
Estimation of lag in analog incorporation. Cells were pulse labeled with 2 μM CldU for 5 minutes and chased with 20 μM IdU for varying lengths of time: 3, 4, 5, 6, 7, 8, 10, 14, 16 minutes. At least 100 forks were collected for each time point except for 3 and 4 minutes of IdU labeling for which only 11 and 44 forks were measured, respectively, due to lack IdU 925 labeled forks at such short lengths of labeling period.

**Figure S14:**
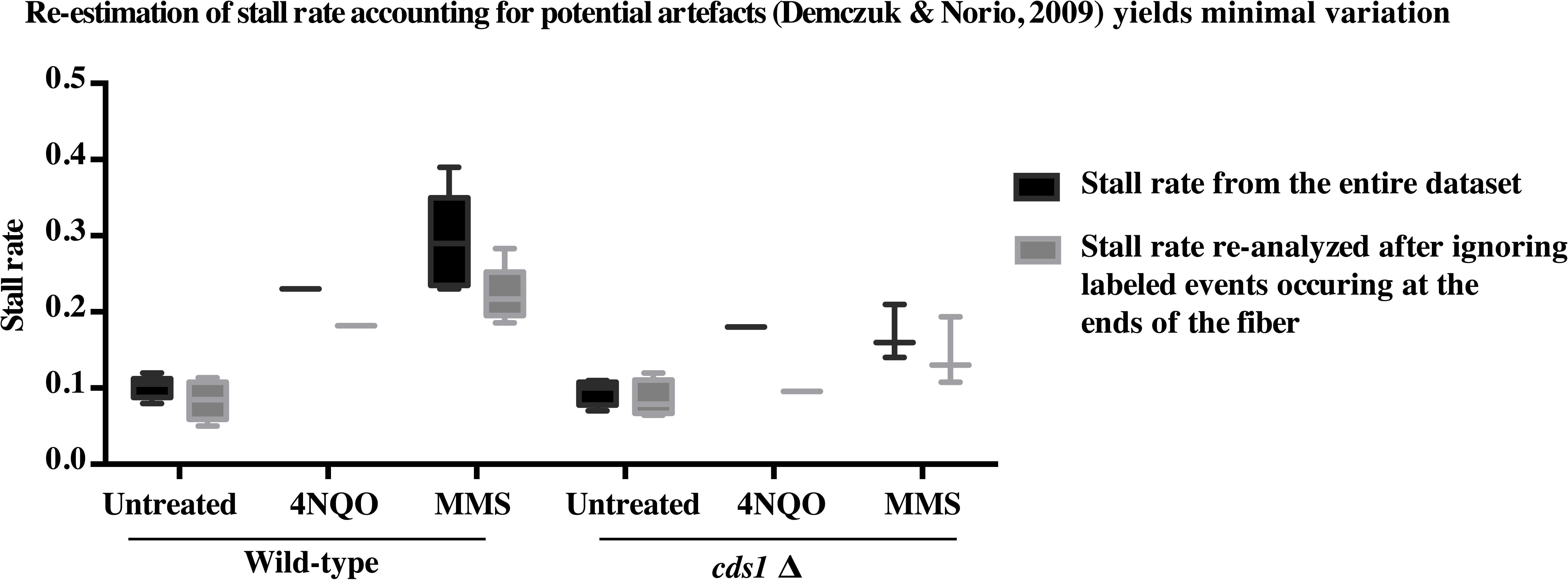
Re-estimation of stall rate accounting for potential artifacts yields minimal variation. We re-analyzed the fibers considering stretching artifacts, which may lead to incorrect interpretation of analog incorporation patterns (Demczuk and Norio, 2009). Ends of the fiber are most susceptible to such stretching artifacts and hence we ignored the labeled events occurring at the ends of the fiber and re-calculated the stall rate. However we do not see a significant change between the stall rate calculated from the whole fiber or after ignoring the labeled events occurring at the ends of the fiber. Wild-type experiment in 4NQO and MMS were done once and 5 times respectively. *cds1*Δ experiment in 4NQO and MMS were done once and thrice respectively.

**Figure S15:**
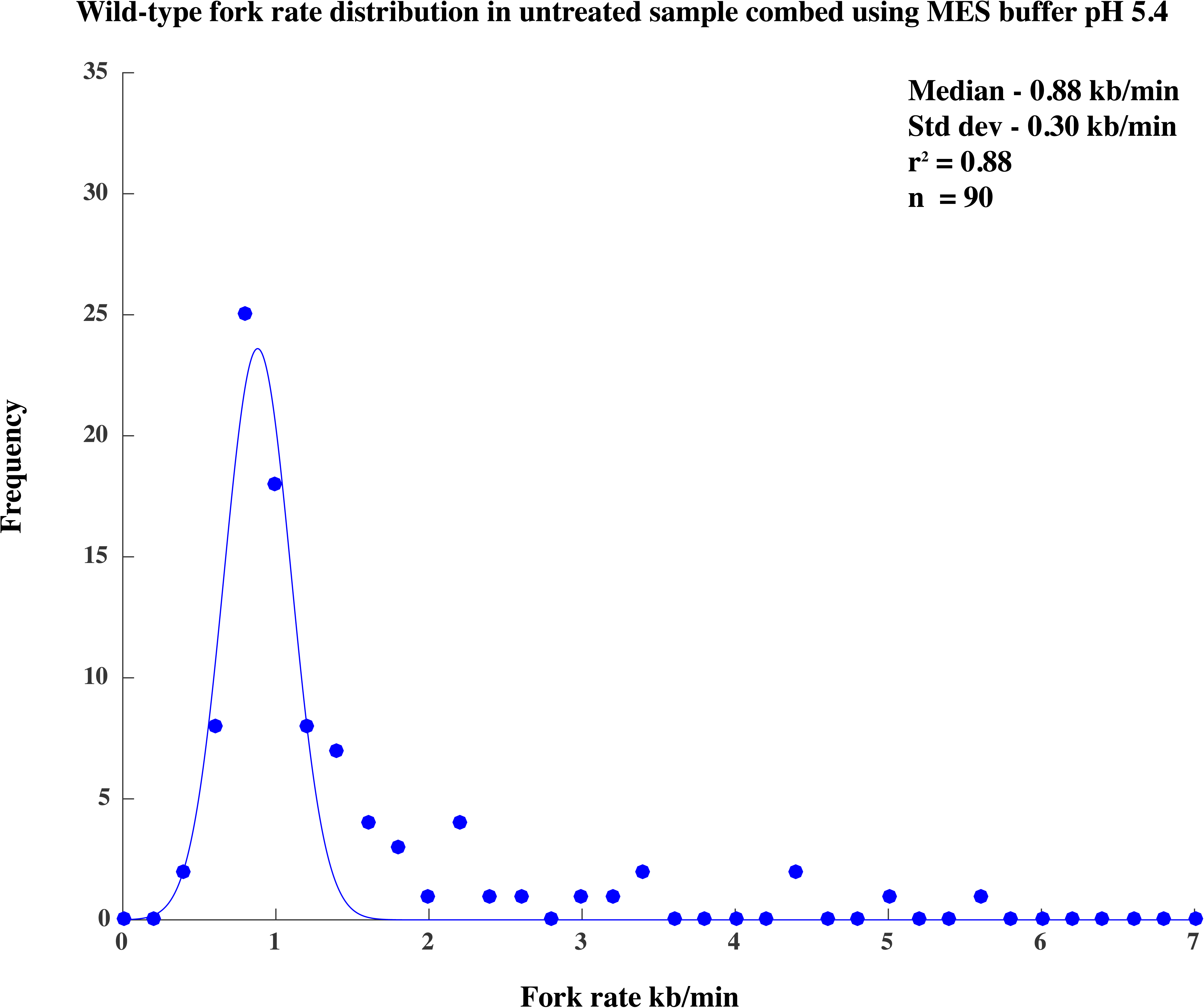
Fork rate distribution in wild-type untreated sample combed using MES buffer pH 5.4. Fork rate value was estimated from the second analog track continuing from the first analog as explained in the Methods section.

**Figure S16:**
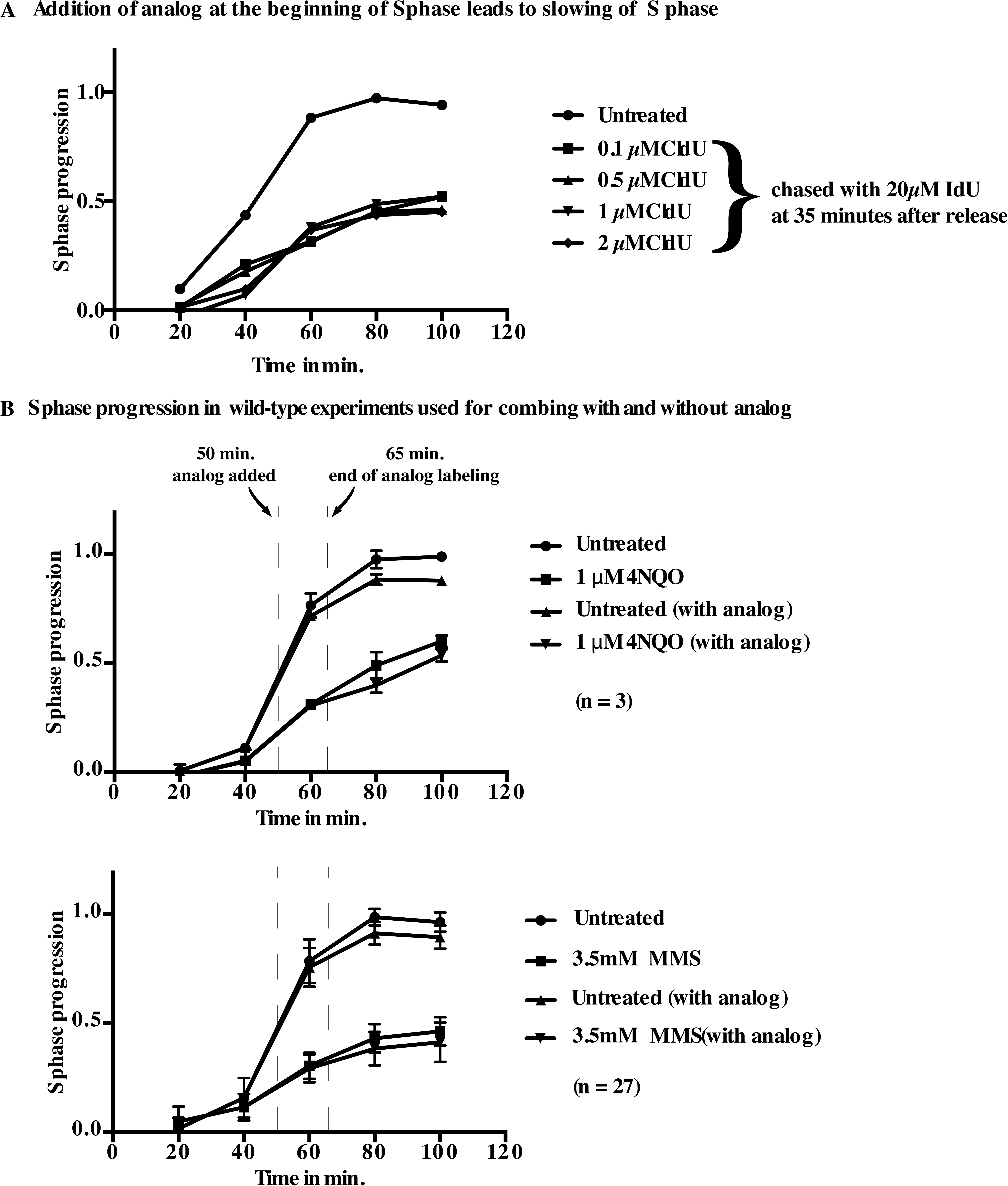
Effect of analog addition on replication kinetics. A) Wild-type cells were synchronized and released into S phase with analog added at different concentrations—0.1 μM, 0.5 μM, 1 μM, 2 μM CldU—and chased with 20 μM IdU at 35 minutes after release or without any analog. B) The S-phase progression of cultures with and without analog addition used for combing experiments. 2 μM CldU analog was added at 50 minutes after release for 5 minutes and chased with 20 μM IdU for 10 minutes and the cells were harvested for combing at 65 minutes after release so as to have minimal effects of analog addition on replication kinetics.

